# Monoaminergic neurons share transcriptional identity across Bilaterian animals

**DOI:** 10.1101/2025.10.10.679534

**Authors:** Matthew Goulty, Clifton Lewis, Filippo Nicolini, Ruman Khalid, Hammad Syed, Paola Oliveri, Mervyn G. Thomas, Ezio Rosato, Roberto Feuda

**Affiliations:** Department of Genetics, Genomics and Cancer Sciences, University of Leicester, Leicester, UK; The Francis Crick Institute, London, UK; Ulverscroft Eye Unit, School of Psychology and Vision Sciences, College of Life Sciences, University of Leicester, Leicester, UK; Centre for Life’s Origins and Evolution, Department of Genetics, Evolution and Environment, University College London, London, UK; Department of Biology, Geology and Environmental Science, University of Bologna, Bologna, Italy

## Abstract

The evolutionary conservation of cell types over deep time has long been theorised but remains difficult to demonstrate. Monoaminergic neurons, which produce molecules such as serotonin and dopamine, are central to animal behaviour and cognition, yet their evolutionary origins remain unresolved. Here, we analysed single-cell transcriptomes from 16 metazoan species spanning eight phyla. Using a novel integration method, Orthogroup Recoding, together with experimental validation, we show that monoaminergic neurons form consistent transcriptional clusters across Bilateria. These clusters share a regulatory signature involving conserved transcription factors, including homologues of *Fev*, *Lmx1b*, *Fer2*, *Insm*, and *Isl*, suggesting a common regulatory program. This signature extends beyond the brain including other monoaminergic cells, such as the gut enterochromaffin cells and larval sensory neurons. By contrast, non-bilaterian lineages lack the coordinated expression of the transcription factors and the biosynthetic machinery that regulate monoamine production. Our observations support the existence of a shared set of TFs that define ‘monoaminergic identity’ across in bilaterians.

## Introduction

The homology of cell types in distant related organisms has long been hypothesised, but for most cell types it remains unresolved^1,2^. Comparisons have typically centred on a few canonical regulatory genes^3^, representing only a fraction of the transcription factors (TFs) that control cell identity. Single-cell transcriptomics (scRNAseq) offers an unprecedented opportunity to comprehensively and unbiasedly compare cell-type expression across species^2,4^. So far scRNAseq comparison of cell types across Metazoa have been limited to broad cell types^5,6^. More specific cell-type comparisons have been limited to few species^7,8^ and/or closely related species^9,10^. Testing conservation of specific cell types across Metazoa requires cell types that are both molecularly well-defined and broadly represented across phyla.

Monoaminergic neurons provide such a system. They are rare (<1% of neurons in vertebrate or insect brains)^11–16^ specialised cells producing serotonin, dopamine, and related molecules that act as key neuromodulators, regulating behaviour and cognition^17–20^. Moreover, they are molecularly defined by a unique enzymatic toolkit that converts amino acids into monoamines, and by transporters that package them for release^21–24^. Monoaminergic cells are found across bilaterians including phyla separated by more than 500 Ma^21,22,25–30^. These cells are found most prominently in the nervous system but also in other tissues, such as the gut^31^.

Current models define cell-identity by the combinatorial expression of transcription factors (TFs)^3,11,32^. Therefore, homologous cell-types should express similar combinations of TFs^1,9,33^. Multiple TFs have been characterised in monoaminergic neurons, primarily in vertebrates and nematodes (Table S1)^11,21,24,34–39,39–71^. However, whether related TFs control a shared ‘monoaminergic identity’ across species is unclear. If this is the case, and monoaminergic neurons represent a genuine homologous feature across species, it would change our understanding of complex behaviours such as learning, arousal, and aggression.

Here, we analyse single-cell RNA-seq data from 16 species^7,11,72–89^ spanning eight phyla to ask whether monoaminergic neurons share conserved transcriptional and regulatory signatures. We used three different methods: (i) orthology-aware alignment (SAMap)^5,90^; (ii) a novel integration method based on hierarchical orthogroup recoding; and (iii) species-specific pseudobulk analyses. We have identified consistent transcriptional similarities in monoaminergic cell types across Bilateria. This includes a common pool of transcription factors associated with monoaminergic identity. We used Hybridization chain reaction (HCR) imaging to validate these predictions in three representative species. Our results reveal a shared transcriptional state and set of TFs which supports a ‘monoaminergic identity’ within Bilateria.

### Monoaminergic neurons form a conserved cross-species transcriptomic cluster

Whether monoaminergic neurons share a conserved transcriptional signature across bilaterian animals remains unresolved. To address this, we applied SAMap^5^, an orthology-aware tool that identifies cross-species cell-type correspondences based on transcriptomic similarity. Unlike methods restricted to 1:1 orthologs, SAMap incorporates many-to-many gene relationships and iteratively refines gene-gene mappings using both sequence similarity and observed co-expression. SAMap computes a similarity score between cell clusters, with values near 1 indicating highly similar transcriptomic profiles and 0 indicating no detectable similarity.

We applied SAMap to brain/neuronal scRNA-seq datasets from 12 bilaterian species (the bilaterian dataset), encompassing major lineages including lophotrochozoans^7^, ecdysozoans^11,81,85–87,91^, and vertebrates^75–77,79,89^ (Fig. 1A, Table S2). Monoaminergic neurons form a well-aligned, cross-species cluster (Fig. 1B–C). Interestingly we did not identify stronger alignment between monoaminergic sub-types (e.g., dopaminergic, serotonergic) across phyla. This suggests conservation of transcriptomic profiles at the broader monoaminergic neuron level, rather than specific subtype homologies. To check whether monoaminergic pathway genes drove the similarities observed in SAMap we calculated the significant gene pairs supporting the monoaminergic cell-type alignments. In addition to monoaminergic pathway genes, SAMap alignments were supported by a variety of homologous genes including synaptic, neuronal, and RNA-processing genes (Table. S3). These results suggest that monoaminergic neurons collectively have a similar expression profile across Bilateria.

**Figure 1:**
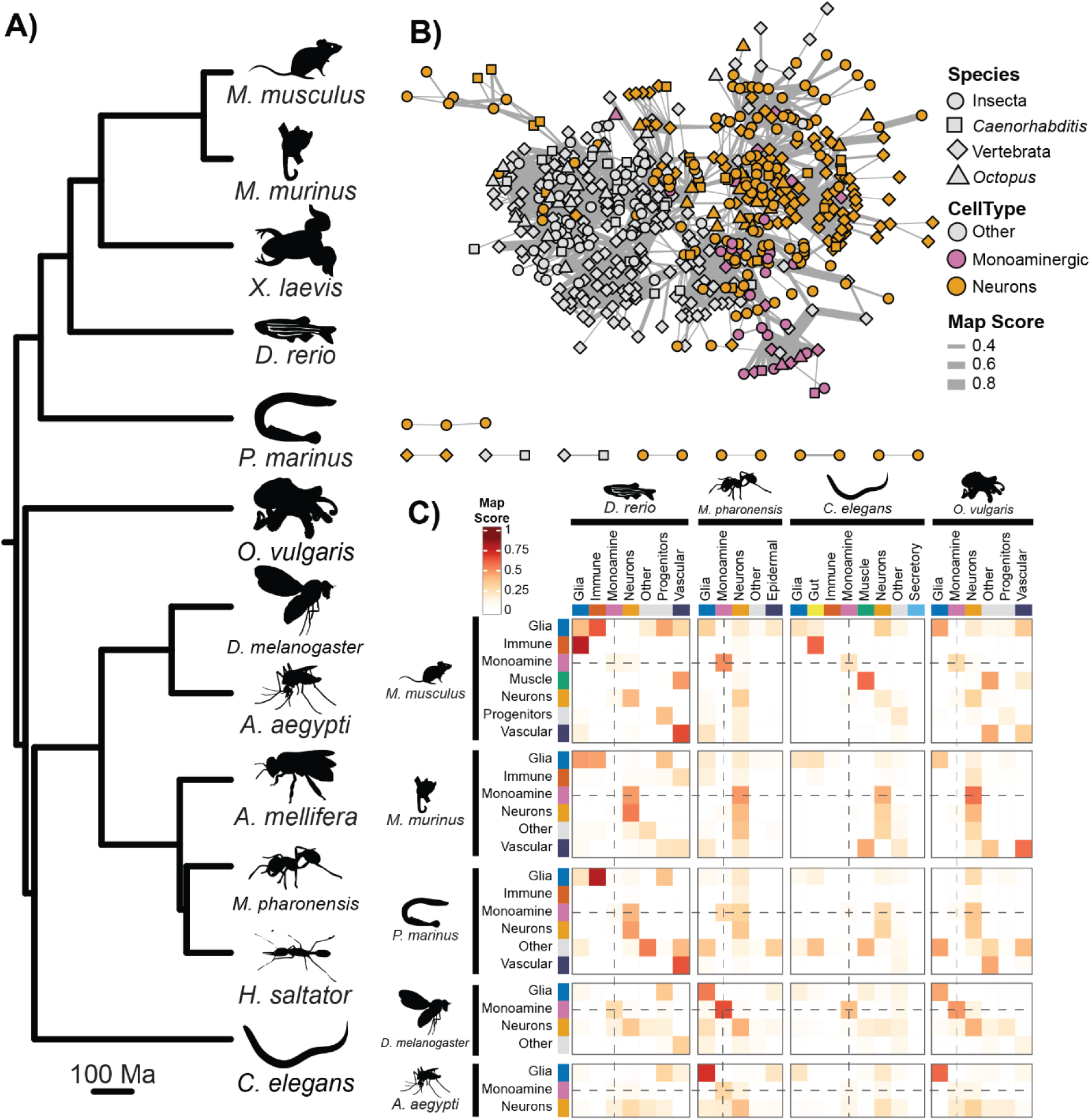
Monoaminergic Neurons share Transcriptional Profiles Across Bilateria. A) Time-scaled phylogeny of species used in the Bilateria brain dataset. B) Network of SAMap mapping scores between individual cell clusters. Nodes represent cell clusters and edges represent SAMap mapping scores. Edge thickness denoted the mapping strength, scores below 0.2 are omitted. C) Heatmap of SAMap mapping scores. Cell clusters have been summarised into broad groupings. Dashed lines follow the row/columns of monoaminergic groupings. Phylogenetic tree dated according to literature^25,108,109,127,128^. Silhouettes from Phylopic.org: silhouette images are by Daniel Jaron (*Mus musculus*), Jake Warner (*Danio rerio*), Margot Michaud (*Octopus vulgaris*), Mattia Menchetti (*Apis mellifera*), Ramiro Morales-Hojas (*Drosophila americana* as *Drosophila melanogaster*), Richard J. Harris (*Aedes albopictus* as *Aedes aegypti*, *Solenopsis invicta* as *Monomorium pharaonis*), Taylor Medwig-Kinney (*Caenorhabditis elegans*), Vijay Karthick (*Harpegnathos saltator*), and others (*Microcebus murinus*, *Petromyzon marinus*, *Xenopus laevis*).

### Orthogroup Recoding directly clusters monoaminergic neurons across phyla and identifies shared transcription factors

To test whether monoaminergic neurons express homologous transcription factors across species, we developed a novel approach to directly compare cross-species expression: Orthogroup Recoding. This method aggregates the expression of all genes within phylogenetically defined orthogroups (at a consistent phylogenetic node), enabling cross-species comparison of genes with complex orthology. We benchmarked Orthogroup Recoding on mammalian pancreas and vertebrate heart datasets based on the BENGAL pipeline^92^ and found that it matched or outperformed standard 1:1 ortholog-based integration across most metrics (Fig. S1-3). Orthogroup Recoding provided the most improvement when comparing species with more complex orthology relationships (across vertebrates) yielding improved cell-type separation and species mixing (Fig. S1-3). This approach allows for direct integration and quantitative comparisons of gene expression across broad evolutionary distances and enables incorporating a more comprehensive set of genes.

Next, using Orthogroup Recoding and Seurat RPCA (Reciprocal Principal Component Analysis), we integrated scRNA-seq data from the bilaterian dataset. Annotated monoaminergic neurons formed four clusters which all displayed clear monoaminergic expression patterns (Fig. 2B). Cluster 19 showed a broad monoaminergic identity expressing all canonical pathway orthogroups (Fig. 2B) and contained monoaminergic neurons from all species (Fig. 2C). Two clusters (23 and 26) consisted almost exclusively (>99%) of *Octopus vulgaris* dopaminergic neurons (Fig, 2C). Cluster 33 only expressed part of the canonical serotonergic pathway (lacking the expression of SLC6 and VMAT orthogroups). Instead, cluster 33 uniquely expressed melatonin biosynthesis genes and predominantly contained vertebrate cells (Fig. 2B).

**Figure 2:**
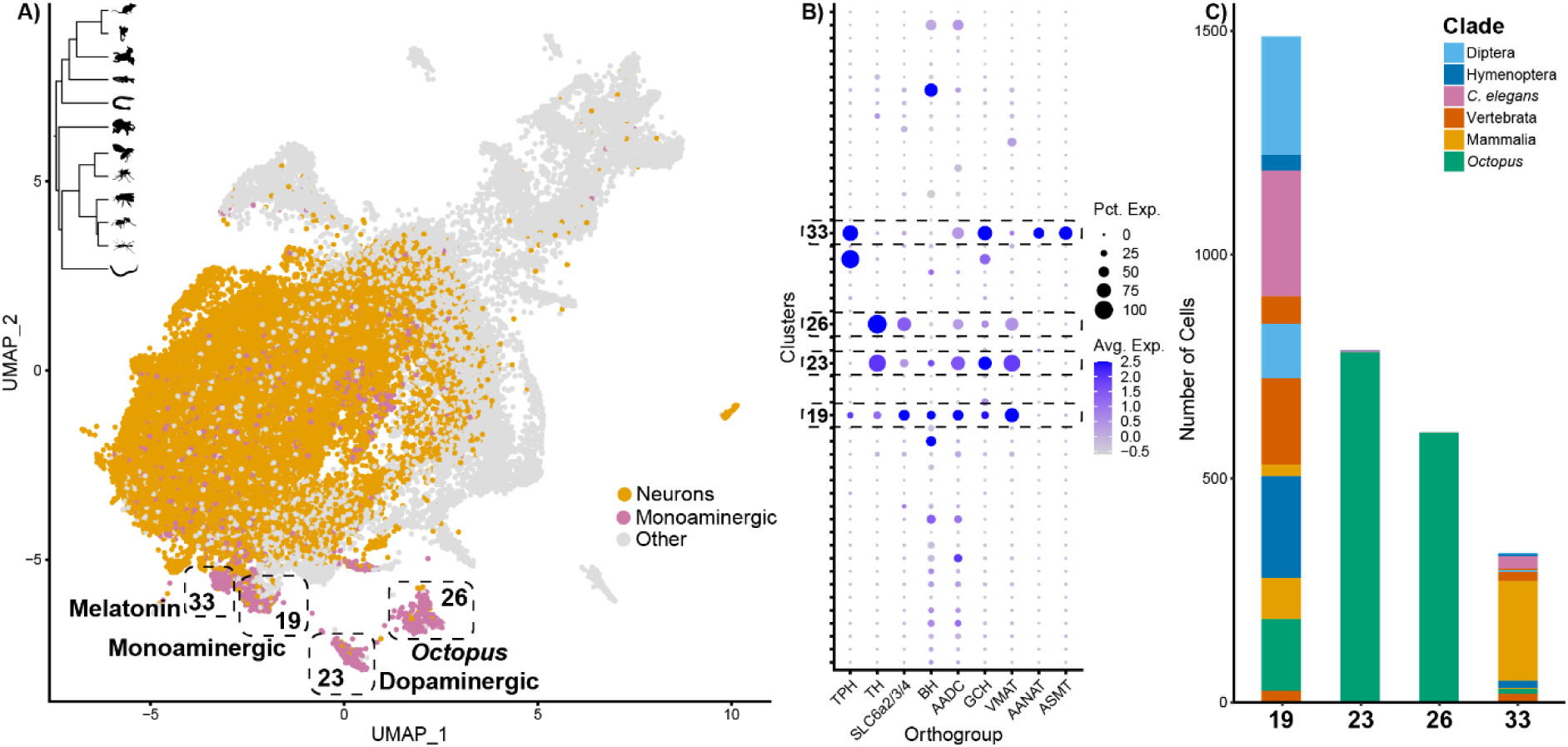
Monoaminergic Neurons Cluster Together. Plots detailing the clustering and expression of monoaminergic cells from 12 brain datasets under Orthogroup Recoding. A) RPCA integrated UMAP, colour coded by annotation. Monoaminergic clusters highlighted with dashed boxes and labelled. Tree of included species inset in top left. B) Dot plot showing the expression of monoaminergic orthogroups across clusters. C) Bar plots of the species makeup of monoaminergic clusters. Gene expression was recoded using the Metazoa level of EggNOG annotations. Silhouettes from Phylopic.org: silhouette images are by Daniel Jaron (Mus musculus), Jake Warner (Danio rerio), Margot Michaud (Octopus vulgaris), Mattia Menchetti (Apis mellifera), Ramiro Morales-Hojas (Drosophila americana as Drosophila melanogaster), Richard J. Harris (Aedes albopictus as Aedes aegypti, Solenopsis invicta as Monomorium pharaonis), Taylor Medwig-Kinney (Caenorhabditis elegans), Vijay Karthick (Harpegnathos saltator), and others (Microcebus murinus, Petromyzon marinus, Xenopus laevis).

**Figure 3:**
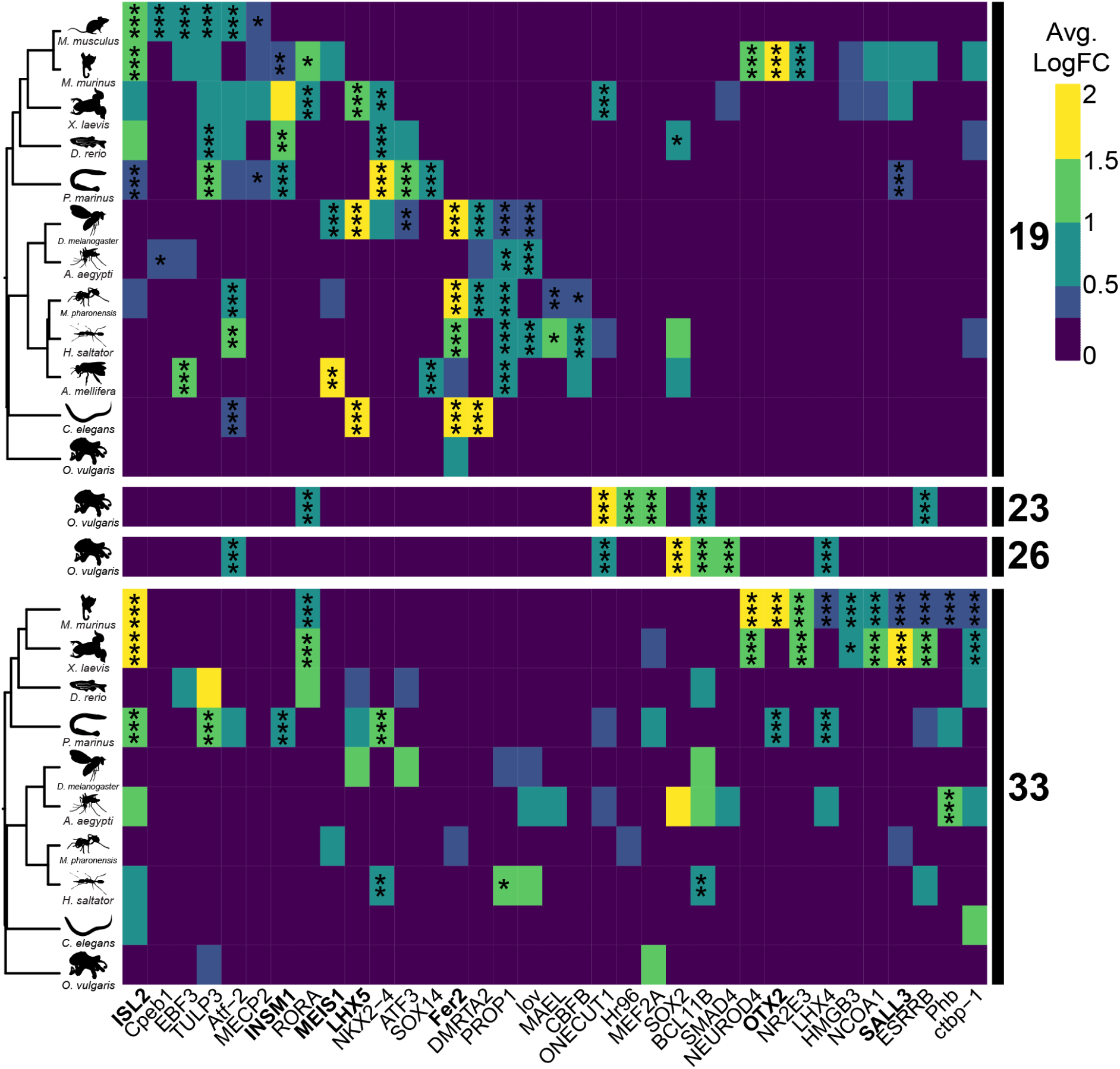
Transcription Factor Orthogroups are Expressed in Monoaminergic Clusters Across Species. Heatmaps of differentially expressed transcription factor orthogroups shared across species in the four monoaminergic clusters. Species trees and icons displayed on the left indicating the species included. Orthogroup names shown along the bottom, orthogroups listed in table 1 are in bold. Orthogroups are included when they are expressed in two or more species in the cluster of interest. For clusters 23 and 26 only orthogroups with > 1 lfc in *Octopus vulgaris* are included. Orthogroups are mapped across all four monoaminergic clusters if shared in one cluster. The colour code corresponds to log fold change expression < 0.25 lfc is omitted, *** indicates p.adj < 0.001, ** indicates p.adk<0.01 and * indicates p.adj<0.05. Silhouettes from Phylopic.org: silhouette images are by Daniel Jaron (*Mus musculus*), Jake Warner (*Danio rerio*), Margot Michaud (*Octopus vulgaris*), Mattia Menchetti (*Apis mellifera*), Ramiro Morales-Hojas (*Drosophila americana* as *Drosophila melanogaster*), Richard J. Harris (*Aedes albopictus* as *Aedes aegypti*, *Solenopsis invicta* as *Monomorium pharaonis*), Taylor Medwig-Kinney (*Caenorhabditis elegans*), Vijay Karthick (*Harpegnathos saltator*), and others (*Microcebus murinus*, *Petromyzon marinus*, *Xenopus laevis*).

Testing the expression of TF orthogroups (Table S4) found no single orthogroup significantly differentially expressed across all 12 species in cluster 19, but strong phylogenetic patterns emerged. Insects and other protostomes shared PROP1, lov, and DMRTA2 orthogroups, while vertebrates consistently expressed ISL, TULP3, and INSM. Other TF orthogroups (Atf-2/NPDC1, MEIS and Fer2) were upregulated in multiple species across phyla. Our results point to the common expression of homologous TFs in monoaminergic neurons, in some cases reaching across phyla.

### Clade-specific regulatory signatures reflect divergent monoaminergic neuron evolution

To investigate how phylogenetic distance affects transcriptional comparisons and regulatory conservation, we created clade-specific datasets for insects and vertebrates. Applying SAMap revealed high alignment scores among insect monoaminergic neurons, which clustered together across species and subtype (Fig. S5). In contrast, monoaminergic neurons in the vertebrate dataset were more heterogeneous (Fig. S6) and strong alignment was seen between melatonin-producing cells (expressing ASMT and AANAT). *Mus musculus* monoaminergic neurons showed poor alignment to other vertebrates. This is despite strong alignment of other cell types (astrocytes, oligodendrocytes etc) (Fig. S6). These discrepancies likely reflect both biological divergence and technical variability and are potentially linked to the rarity of monoaminergic neurons in the brain.

In insects, integration using Orthogroup Recoding revealed a single monoaminergic cluster (cluster 18) comprising cells from all five species and enriched for monoaminergic pathway orthogroups (Fig. S10). Eight TF orthogroups were upregulated in two or more species (Fig. S10). These included CG32523/PROP1 (enriched in all five species), Fer2, MEIS1, and DMRTA2. In vertebrates, Orthogroup Recoding again recovered a clear melatonin cell type (cluster 28) where uniquely NR2E3, LHX4 and SALL1/3 orthogroups were enriched in three out of four species (S11). Other monoaminergic neurons formed subclusters within broad neuronal clusters. Subclusters 3 and 1 showed broad monoaminergic and dopaminergic identities respectively. The subclusters expressed distinct but overlapping TF profiles, including DLX5, TULP3, and MEIS. Interestingly ISL was differentially expressed in both subcluster 3 and the melatonin cell type.

### Monoaminergic transcriptional profiles span tissues and life stages

Monoaminergic cells appear in diverse anatomical and developmental contexts, such as apical sensory organs in marine larvae^80,93,94^, arthropod ventral nerve cords^13,21^ and enterochromaffin cells in vertebrate guts^31,95^. To test whether these diverse cell types share core transcriptional features, we integrated a mix of neuronal^11,76,79,81,83,84,89^, gut^74,76,82^, and whole-body scRNA-seq^78,80^ data from six model organisms. Despite varying sample types, monoaminergic cells clustered together (cluster 20), including neurons from *D. melanogaster* ventral nerve cord, mammalian enterochromaffin cells, and monoaminergic cells from *Strongylocentrotus purpuratus* larvae. A second cluster (41), composed exclusively of *M. murinus* and *D. rerio* cells, expressed melatonin pathway genes. In cluster 20 many TFs orthogroups were differentially expressed across taxa. Atf-2/NPDC and FEV were expressed in all six species; LMX1 was found in several; INSM and ISL were shared between vertebrates and *S. purpuratus*; and PROP1, Fer2, and LHX5 were shared by *D. melanogaster* and *C. elegans*. These findings suggest links between monoaminergic pathway expression across different tissues and developmental stages.

### Pseudobulk analysis confirms conserved transcriptional patterns

To validate that TF expression patterns were not methodological artefacts of integration or Orthogroup Recoding, we performed differential expression analyses of transcription factors on individual single-cell datasets (Fig. 5A,S18-23). For datasets containing >=3 biological samples we pseudo-bulked the data by sample to mitigate sampling/batch effects.

**Figure 4:**
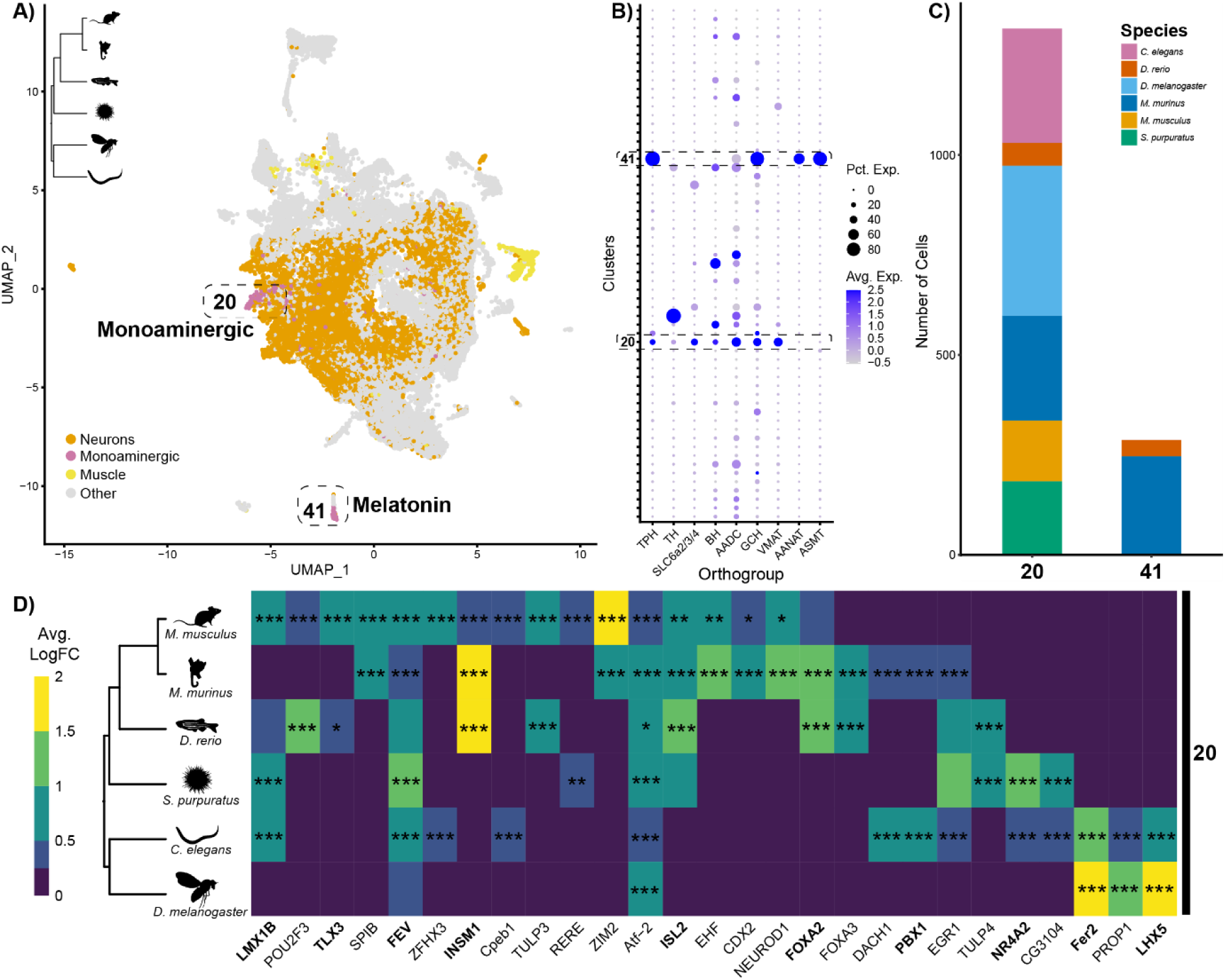
Brain, Larval and Gut Monoaminergic Cells Share Transcription Factor Profiles. Plots detailing the clustering and expression of monoaminergic cells from diverse samples and model organisms integrated under Orthogroup Recoding. A) RPCA integrated UMAP colour coded by annotation. Monoaminergic clusters highlighted by a dashed box and labelled. A phylogenetic tree of included species inset in the top left. B) Dot plot showing the expression of monoaminergic orthogroups across clusters. C) Bar plots showing the species make up of monoaminergic clusters 20 and 41. D) Heatmap of differentially expressed transcription factor orthogroups shared across species in cluster 20. Species tree and icons displayed on the left. Only orthogroups significantly expressed in >= 2 species included. Colour coded by log fold change expression, *** indicates p.adj < 0.001, ** indicates p.adk<0.01 and * indicates p.adj<0.05. Phylogenetic tree dated according to literature^25,108,109,127,128^. Silhouettes from Phylopic.org: silhouette images are by Christoph Schomburg (*Strongylocentrotus purpuratus*), Daniel Jaron (*Mus musculus*), Jake Warner (*Danio rerio*), Ramiro Morales-Hojas (*Drosophila americana* as *Drosophila melanogaster*), Taylor Medwig-Kinney (*Caenorhabditis elegans*), and others (*Microcebus murinus*).

**Figure 5:**
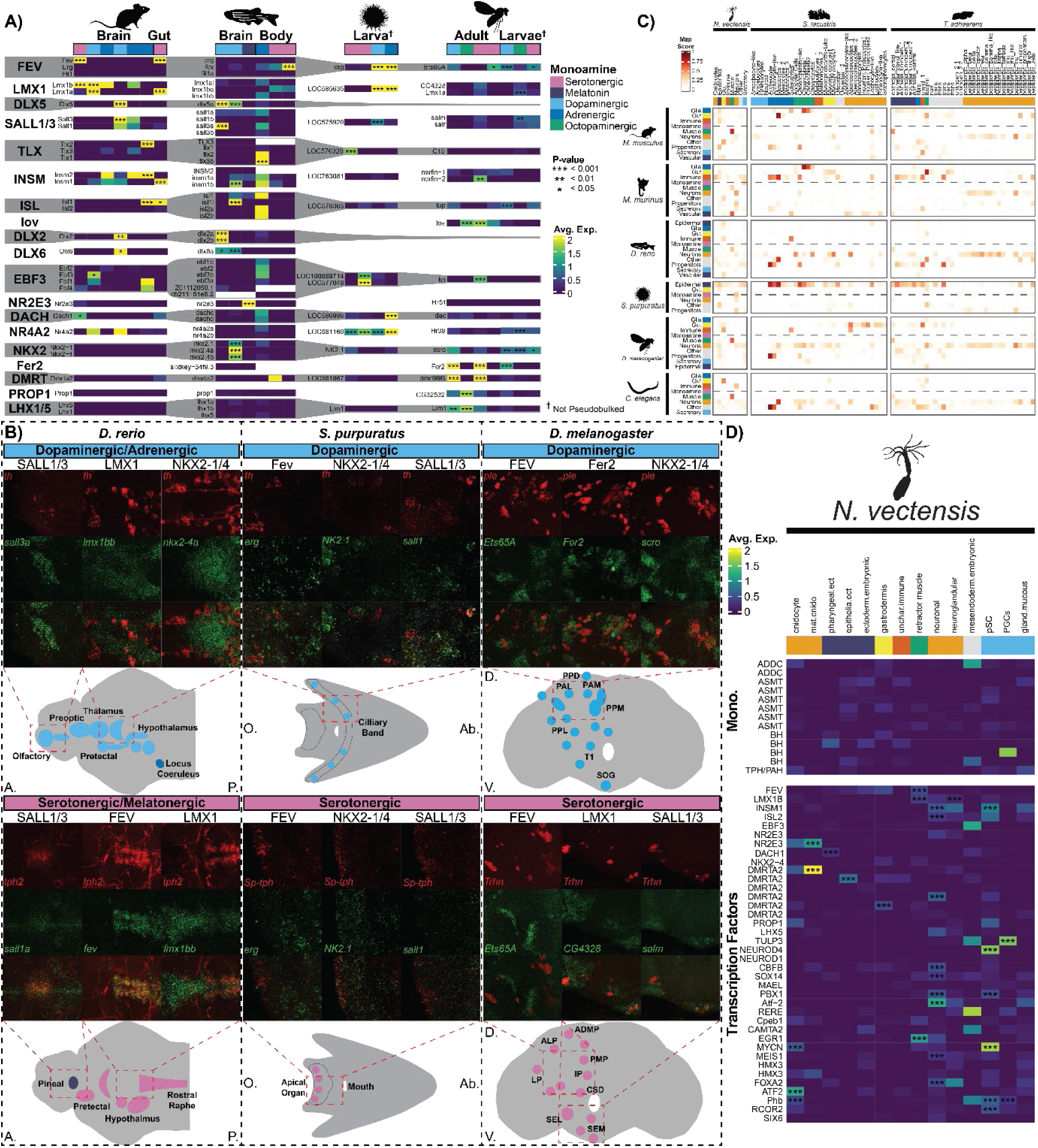
Bilaterians Co-express Common Transcription Factors and Monoaminergic Genes while non-bilaterians do not. A) Linked heatmaps of transcription factor expression from pseudobulk analyses. Grey and white regions indicate orthogroups with individual homolog expression show for each species. Monoaminergic clusters shown with monoamine subtype colour coded along the top. Black boxes and labels group clusters from the same dataset. Significance of differential expression. Indicated by *. Non-pseudo-bulked datasets indicated with †. Orthogroup names shown along the right. Not all significant orthogroups shown, full figures for each species are found in supplement. B) Fluorescent HCR *In Situ* images showing co-expression of selected transcription factors and monoaminergic biosynthetic enzymes. Dashed boxes group images by species with co-expression against dopaminergic enzymes (*th2*, *th*, *ple*) on the top and co-expression with serotonergic enzymes (*tph1a*, *tph2*, *Sp-Tph*, *Trhn*) along the bottom. Grey schematics of the whole samples shown with known monoaminergic cell types indicated and labelled based on the literature^12,13,22,48,129,130^. *S. purpuratus* schematic based on Slota and McClay, 2018^48^. Red dashed boxes highlight the sections of the sample displayed in the fluorescent image. All enzymes coloured in red and transcription factors coloured in green. Fluorophores used are: Alexa-488, Alexa-594, Alexa-546 and DAPI (shown in supplement). Gene and probe information detailed in Table S5. C) Heatmaps of SAMap mapping scores showing alignment between bilaterian and non-bilaterian species. Non-bilaterian species shown as columns and bilaterian species shown as rows. Dashed lines follow monoaminergic rows. D) Heatmap of monoaminergic pathway and transcription factor gene expression in *Nematostella vectensis*. Significance of differential expression is indicated by * Cell type annotations colour coded by broad grouping. Silhouettes from Phylopic.org: silhouette images are by Christoph Schomburg (*Strongylocentrotus purpuratus*), Daniel Jaron (*Mus musculus*), Jake Warner (*Danio rerio*, *Nematostella vectensis*), Ramiro Morales-Hojas (*Drosophila americana* as *Drosophila melanogaster*), Yan Wong (*Trichoplax adhaerens*), and others (*Isodictya grandis* as *Spongilla lacustris*).

In *M. musculus*, homologs of FEV, LMX1, and INSM marked multiple monoaminergic populations, including midbrain and hindbrain neurons and enterochromaffin cells. Olfactory dopaminergic neurons expressed SALL1/3, DLX1, and DLX2. NR2E3 and LHX4 marked melatonin-producing neurons across vertebrates. In *D. melanogaster,* adult brain monoaminergic neurons expressed PROP1, lov, and DMRTA2. Larval neurons from *D. melanogaster*, however, expressed vertebrate-like TFs such as *tup* (ISL2), and *scro* (NKX2.4). *Fer2*, both LMX1 homologs and *Ets65A* (FEV) were shared across developmental stages. These results confirm that conserved TFs are biologically relevant and that between species TFs are differentially used across life stages and monoaminergic subtypes.

### *In Situ* mRNA Mapping Verified Co-expression Patterns

For selected TFs (Table S5) we mapped their co-expression with monoamine biosynthetic enzymes using fluorescent *in situ* HCR (Fig 5B).

In the *D. rerio* developing brain we found clear patterns of expression matching previous publications and the scRNAseq data^21,96,97^. There was strong co-expression of *fev*, *lmx1bb* and *tph2* in the raphe nuclei as identified elsewhere^62,98^. There was limited evidence of this in the scRNAseq data, likely reflecting the insufficient sampling of rare cell types^14^. *Nr2e3* clearly marked the pineal region with highly specific expression (and co-expression with *tph2*) indicating this as the source of the melatonin synthetic cells. *Sall1a was* also expressed there. The midbrain dopaminergic neurons showed co-expression between *lmx1bb*, *nkx2.4a* and *th*. *Sall3a* and *th* co-expression was found in the dopaminergic neurons of the olfactory bulb (Fig 5B and S15).

The analysis of *S. purpuratus* larvae matched some of our expectations. There was clear co-expression of *NK2.1* with *Sp-Tph* but not with *th*^99^. Conversely, we found expression of *sall1* in some *th+* cells while none in *Sp-Tph+* cells, which matched the scRNAseq data. *Erg* did show a clear expression in the skeletal filaments (identified in the scRNAseq) and did not co-locate with *Sp-Tph* as expected. However, the co-expression of *erg* with *th* was weak, contradicting the scRNAseq where *erg* was more upregulated in the dopaminergic neurons than in skeletal cells (Fig 5B and S16).

In adult *D. melanogaster* brains, the transcription factors consistently co-expressed with the enzymes, even in cases where we observed non-significant upregulation in the pseudobulk analyses. For all TFs co-expression was non-exclusive. Both TFs and enzymes were expressed in multiple regions with partial overlap. The co-expression between *CG4328* and *salm* with *Trhn* mapped to the ventral serotonergic neurons while *Ets65A* co-expressed with *Trhn* elsewhere, matching the two serotonergic clusters in the scRNAseq data (Fig 5B and S17). *Ets65A*, *Fer2*^100^ and *scro* all co-expressed with populations of *ple+* neurons in the dorsal region of the brain (Fig 5B and S17).

Overall, HCR expression mapping confirms the general picture derived from the single-cell and is in agreement with published results. However, it has highlighted a few key caveats. Firstly, single-cell data only samples gene expression. This leads to cell groupings which approximate cell type definitions, inevitably missing the fine resolution heterogeneity of specific identities that are present in reality (especially for rare cell types). Secondly, although we assume that the expression levels in scRNAseq are roughly proportional to the level of mRNA in cells, there are collection, amplification and sequencing biases that can distort the pattern. Moreover, post-transcriptional factors are omitted from scRNAseq data. Thus, it is essential to validate computational inferences *in situ*.

### Non-bilaterians lack coordinated monoaminergic transcriptional programs

Monoamine responses have been reported in non-bilaterian metazoans^101,102^. However, while monoaminergic signalling components exist in non-bilaterians, the majority of the enzymatic machinery appears to be bilaterian-specific^103^. Therefore, the presence of monoaminergic neurons in non-bilaterian animals is unclear. We investigated this using available scRNA-seq data from *Nematostella vectensis*^73,104^, *Trichoplax adhaerens*^88^, and *Spongilla lacustris*^72^. Initially we compared the non-bilaterians with the model species scRNAseq dataset using SAMap and we observed a low level of similarity between monoaminergic neurons and the neurons of *Nematostella vectensis* (Fig 5C). However, there was no strong or consistent mapping of non-bilaterian clusters with specific monoaminergic clusters (Fig S24). Next, we investigated the expression of homologs of monoaminergic pathway genes and TFs of interest (Fig 5B, S12-14). Our results suggest that despite possessing homologs of monoaminergic pathway genes and TFs, non-bilaterians do not show consistent co-expression of monoaminergic pathway genes nor their coherent co-expression with key transcription factors (Fig. 5D).

## Discussion

Monoaminergic neurons are fundamental regulators of neuromodulation in bilaterians^17,103,105–107^, yet their evolutionary origin has remained unresolved. By integrating single-cell transcriptomes from 16 metazoan species and validating predictions with HCR, we show that monoaminergic neurons share a common pool of TFs across Bilateria. This signature extends beyond the brain encompassing gut enterochromaffin cells and larval sensory neurons. This suggests that a shared ‘monoaminergic identity’ does reflect biological reality across Bilateria.

We found transcriptionally distinct subtypes that represent lineage-specific innovations. Vertebrate melatonin-producing cells express *ASMT* and *AANAT* together with a unique complement of TFs (OTX2, NR2E3 and LHX4), while cephalopod dopaminergic neurons form separate clusters from other monoaminergic populations. These findings illustrate that cellular identity can be remodelled by lineage-specific pressures to generate novel cell identities.

Patterns of conservation are not explained simply by phylogenetic distance. Insects, despite hundreds of millions of years of divergence^108^, show highly conserved monoaminergic neurons characterised by co-expression of Fer2, DMRT2A, PROP1, lov, and LHX5 homologs. In contrast, vertebrate monoaminergic neurons, despite the similar divergence times^25,109^, display greater heterogeneity and less consistent transcriptional similarity among monoaminergic neurons. These results likely reflect both biological diversification of vertebrate subtypes and the technical challenges of accurately sampling rare cell types.

Our results provide biological support for a shared ‘monoaminergic identity’. Following current models of cell-type evolution, the shared TFs we find may reflect the signature of a homologous ancestral cell type. This would explain TF found in multiple species monoaminergic cells as conserved and infer differences in TF expression as lineage-specific features. However, the different combinations of TFs observed in each species/cell type could mark indirect homology, where lineages have independently converged on using a shared set of TFs. In this scenario lineages independently re-tool the limited number of ancestral regulatory connections controlling monoaminergic pathway genes resulting in similar cell-type signatures.

Overall, our results show that monoaminergic neurons share transcriptional similarity across Bilateria. Indeed, we find a common set of TFs, such as *FEV, Fer2, ISL,* and *INSM*. Importantly, the co-expression of both the TFs and the enzymatic machinery appears to be restricted to bilaterians only, suggesting that the monoaminergic cell identity originated in the bilaterian stem lineage. While broader sampling of non-bilaterian metazoans will be required to draw firm conclusions, our results suggest that responses to monoamines in these lineages^101,102^ likely involve analogous rather than homologous systems.

It remains unclear whether and to what degree the expression of TFs we observe in bilaterians monoaminergic neurons reflects the wiring of a conserved gene regulatory network or the emergence of alternative network architectures that converged on similar transcriptional states. Furthermore, it remains unknown the extent to which monoaminergic neurons are the product of homologous developmental trajectories/processes. Functional tests will be required to distinguish these alternatives. Future studies with more homogeneous sampling and improved single-cell technologies will be needed to identify at greater resolutions the TF combinations that define specific monoaminergic subtypes.

Our work raises new questions about how monoaminergic neurons originated and evolved, and provides initial evidence that cell-type transcriptional programs could be conserved over deep evolutionary time.

## Methods

### Single-Cell Pre-processing

Single-cell RNAseq datasets and their corresponding proteomes were obtained from the literature, (Table S2). These datasets were re-analysed using Seurat v4.4.0^110,111^ (Table S6) to re-cluster and annotate cell types based on markers listed in the source papers as well as markers from https://panglaodb.se/^112^ and neuronal markers identified from literature^74,81,83,89,112–114^.

Monoaminergic pathway genes were identified using EggNOG v5^115^ and EggNOG mapper v2^116^ to annotate the single-cell RNAseq datasets respective proteomes. Expression of these genes were mapped and used to identify monoaminergic clusters.

### SAMap Integration

Single-cell datasets were converted from Seurat to h5ad objects using sceasy^117^. Diamond^118^ was used to run pairwise all-vs-all BLASTP comparisons of species proteomes. SAMap^5,90^ was used to align bilaterian, vertebrate, insect and bilaterian plus non-bilaterian datasets to compare the alignment of monoaminergic cell clusters. Seurat clusters were used for the neighbourhood comparisons and mapping score calculations. Cell type annotations were binned into broader categories and mapping scores were calculated. The pairwise comparison mode of SAMap was used with a BLASTP evalue threshold of 1e-6.

Mapping scores were extracted and used to generate network plots in R^119^ using igraph and ggraph.

### Orthogroup Annotation

All proteomes were annotated with EggNOG V5^115^ using EggNOG-mapper^116^. For all datasets used in Orthogroup Recoding we constructed a master annotation table which matched the feature names from the single-cell expression matrix with their corresponding EggNOG Orthogroup at key phylogenetic levels. Unannotated feature names were kept unchanged.

### Transcription Factor Orthogroup Curation

Verified lists of TFs were downloaded for *H. sapiens*, *M. musculus*, *D. rerio, D. melanogaster* and *C. elegans* from AnimalTFDB v4^120^. These names were cross-referenced with the EggNOG annotations of the respective species proteomes. The annotations were filtered and combined to produce a list of 1,944 EggNOG orthogroups (at the Metazoa level) that contained a verified TF. Orthogroups were named according to constituent genes. (Table S4)

### Benchmarking Analyses

We obtained pancreas and heart datasets listed in the BENGAL paper^92^. These were re-analysed in Seurat and cluster annotations harmonised to match those listed in the scOnto outputs in the supplementary material of Song *et al*^92^.

The homology mapping script from Song *et al.* was modified to identify the ENSMBL 1:1 orthogroups and to perform Orthogroup Recoding by summing the counts for all features annotated to the same EggNOG orthogroup. Recoding was done at the Mammalia, Vertebrata and Metazoa levels (as appropriate). The filtered and recoded datasets were then concatenated across species. We used Song *et al*’s scripts to analyse each dataset using seven integration tools: scANVI, scVI, Seurat CCA, Seurat RPCA, FastMNN, Harmony^121^ and Scanorama^122^. SAMap was omitted as it is not compatible with orthogroup recoding.

We modified the metric calculation and comparison scripts from Song *et al* to generate the scIB^123^ and integration scores. Some metrics were omitted: ACLs as the custom package did not work; the kBET metric from scIB did not run and was removed as was the iFL score from the heart dataset. We altered the integration score weightings correspondingly.

### Seurat Integration

Original count matrices were loaded into R. Datasets were recoded using the master feature tables to group features by their Metazoa level EggNOG annotations. In each cell the counts from grouped features were summed producing cell-by-orthogroup matrices. Recoded matrices were re-analysed using Seurat and annotated as the original data.

Recoded datasets were integrated using the RPCA algorithm. Datasets were subsampled to 10,000 cells, if larger. Highly variable orthogroups were calculated for each dataset and compared to select up to 2000 shared highly variable features (orthogroups). These were used to calculate a PCA for each dataset. The RPCA algorithm compared the PCs between datasets to identify similar cells (anchors). These anchors are used to align the PCA spaces across species and transform the original highly variable features. The transformed integrated values are then used to perform a standard single-cell analyses combining all the datasets.

### Differential Gene Expression Testing

The recoded integrated datasets were split by species. FindAllMarkers was used to test the differential expression of orthogroups in monoaminergic clusters identified in the integrated analysis. Only the 1,944 TF orthogroups were tested (Table S4) and only positively differentially expressed results tested.

### Pseudobulk Differential Gene Expression Testing

The original un-recoded matrices for each dataset were annotated with the biological sample ID. UMI counts were aggregated by sample and Seurat cluster to produce pseudobulk datasets. When datasets contained 3 or fewer samples the original count matrices were kept.

The datasets were tested for marker TFs using FindAllMarkers. Only genes annotated to one of the 1,944 TF Orthogroups and calculated as positively expressed markers were tested. The results were filtered to only markers of the monoaminergic clusters. Results were further filtered to only genes annotated to the orthogroups identified as significantly differentially expressed in 2 or more species in the integrated monoaminergic clusters.

### *In Situ* Hybridisation Chain Reaction

TF orthologs of interest were identified from the pseudobulk expression tests from *Danio rerio*, *Drosophila melanogaster* and *Strongylocentrotus purpuratus*. The mRNA for selected genes was obtained from Flybase^124^, Echinobase^125^ or ENSMBL. Isoforms were aligned in seaview and the longest sequence selected. Selected mRNA sequences were used to design HCR probes (by Molecular Instruments, Table S5). Probes were given adaptors so any enzyme probe could be multiplexed with any transcription factor probe.

Oregon-R WT *D. melanogaster* (Bloomington: 2376) were maintained on standard cornmeal fly food in vials under routine laboratory conditions of light and humidity. Adult flies (∼10–30 days old) were fixed overnight on a nutating mixer in PBS containing 3.7% (v/v) formaldehyde and 5% (v/v) DMSO. Following fixation, flies were washed thoroughly with PBT (PBS + 0.1% Tween-20) to remove residual fixative. Brains were then dissected under a stereomicroscope and processed for HCR staining.

Adult *Strongylocentrotus purpuratus* were obtained from the Monterey Abalone Company and maintained in aquaria at 12 °C. Pluteus larvae were collected at 5 days post-fertilization from cultures reared at 12 °C. Larvae were fixed overnight at 4 °C in 4% paraformaldehyde (Electron Microscopy Sciences, Cat. #15710) in artificial seawater. Following fixation, samples were washed three times in washing buffer (0.1 M MOPS, 0.5 M NaCl, 2 mM EDTA, 1 mM MgCl₂, 1× PBS) and subsequently dehydrated in 100% methanol. Samples were stored long-term at −20 °C.

Adult zebrafish (*Danio rerio*) were maintained in the Preclinical Research facility at the University of Leicester under standard conditions. Breeding was initiated by separating adult males and females overnight with a divider. Fertilised eggs were collected the following morning after removal of the divider. Embryos were maintained in 10-cm Petri dishes containing fish water (0.3 g/L Instant Ocean) at 28.5°C until 5 days post-fertilisation (dpf). Embryos were treated with 200uM PTU (Phenylthiourea) starting from 24 hours post fertilisation and the solution was changed every 24 hours. At 5dpf the embryos were fixed in 4%PFA overnight at 4**°**C. The subsequent steps were performed according to the protocol from Molecular Instruments with minor adjustments.

All procedures adhered to the Animals in Scientific Procedures Act 1986 and were conducted under project licence PP1567795 by researchers holding individual UK Home Office personal licenses.

All samples were processed according to the HCR protocols provided by Molecular Instruments. Pairs of probes were used to map the co-expression of monoaminergic biosynthetic enzymes and potentially conserved transcription factor homologs. Stained samples were imaged using Zeiss LSM 980 Airyscan 2 (AIF, Leicester). The images were processed on Zeiss Zen Blue and FIJI^126^.

Key reagents list in Table S7.

## Acknowledgements

This work is supported by a University Research Fellowship (UF160226 and URF/R/221011) and a Research Grant (RGF\R1\181012), awarded to R.F. M.G. is supported by a PhD studentship from the College of Life Sciences. C.L. is supported by a BBSRC-MIBTP PhD Studentship (Grant number MIBTP2020: BB/T00746X/1). The authors wish to thank the technical and scientific staff of several core facilities for their invaluable support. We are especially grateful for the assistance received from the University of Leicester Advanced Imaging Facility (RRID: SCR_020967; BBSRC grant: BB/S019510/1) and specifically thank Dr. Kees Straatman for his expert microscopy advice. This research used the ALICE High Performance Computing Facility at the University of Leicester.

## Author Contributions

M.G and R.F. conceived of the project. M.G. performed all computational analyses. F.N and R.K performed and imaged HCR on *D. rerio* and *S. purpuratus*. C. L. and R.K performed and imaged HCR on *D. melanogaster*. C.L. maintained, fixed and dissected *D. melanogaster* samples. P.O. provided *S. purpuratus* larval culture and fixed samples. M.T. and H.S. provided *D. rerio* culture and fixed samples. R.F supervised the project. M.G. and R.F edited the manuscript.

## Data Availability

Scripts, metadata and results supporting this study are available via Figshare repository (doi: 10.25392/leicester.data.30217852). Sources of original data used listed in supplementary Table S1. Reanalysed datasets available from authors by request.

## Notes

### Competing Interest Statement

The authors have declared no competing interest.

