## Supplementary Figures for "Monoaminergic neurons share transcriptional identity across Bilaterian animals"

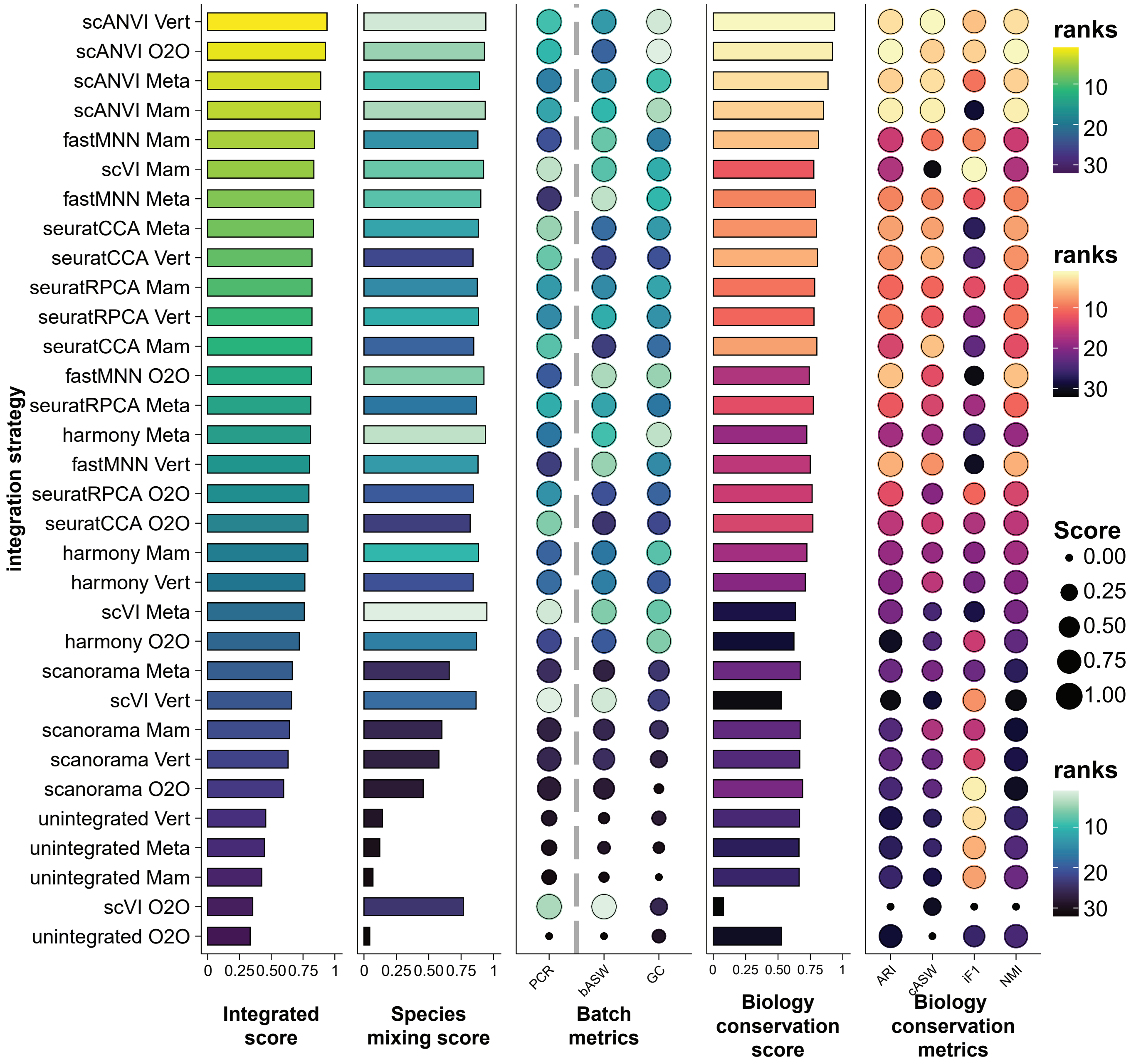

integration strategy
scANVI Vert
scANVI O2O
scANVI Meta
scANVI Mam
fastMNN Mam
scVI Mam
fastMNN Meta
seuratCCA Meta
seuratCCA Vert
seuratRPCA Mam
seuratRPCA Vert
seuratCCA Mam
fastMNN O2O
seuratRPCA Meta
harmony Meta
fastMNN Vert
seuratRPCA O2O
seuratCCA O2O
harmony Mam
harmony Vert
scVI Meta
harmony O2O
scanorama Meta
scVI Vert
scanorama Mam
scanorama Vert
scanorama O2O
unintegrated Vert
unintegrated Meta
unintegrated Mam
scVI O2O
unintegrated O2O
0 0.25 0.50 0.75 1
Integrated score
0 0.25 0.50 0.75 1
Species mixing score
PCR
bASW
GC
Batch metrics
0 0.25 0.50 0.75 1
Biology conservation score
ARI
cASW
iF1
NMI
Biology conservation metrics
ranks
10
20
30
ranks
10
20
30
Score
0.00
0.25
0.50
0.75
1.00
ranks
10
20
30

**Figure S1: Mammal Pancreas Dataset Integration Comparison Scores**

Plots showing the breakdown of benchmarking scores for the integration of human and mouse pancreas datasets. Integration method (tool+orthology method) shown on the left, ordered by overall integrated score. Metrics left to right: Integrated score, overall score averaging the species mixing and biological conservation scores; Species mixing score, an average of the metrics measuring the mixing of cells across species; Batch metrics, three metrics (PCR, bASW and GC) that denote if cells group by batch (in this case species); Biology conservation score, an average of metrics measuring the grouping of cells by their given cell type annotation; Biological conservation metrics, four metrics (ARI, cASW, iF1 and NMI) that make up the Biological conservation score. The orthology methods are abbreviated as follows: O2O = 1:1 orthologs only; Mam = orthogroup recoding at the Mammalia EggNOG level; Vert = orthogroup recoding at the Vertebrata EggNOG level; Meta = orthogroup recoding at the Metazoa EggNOG level.

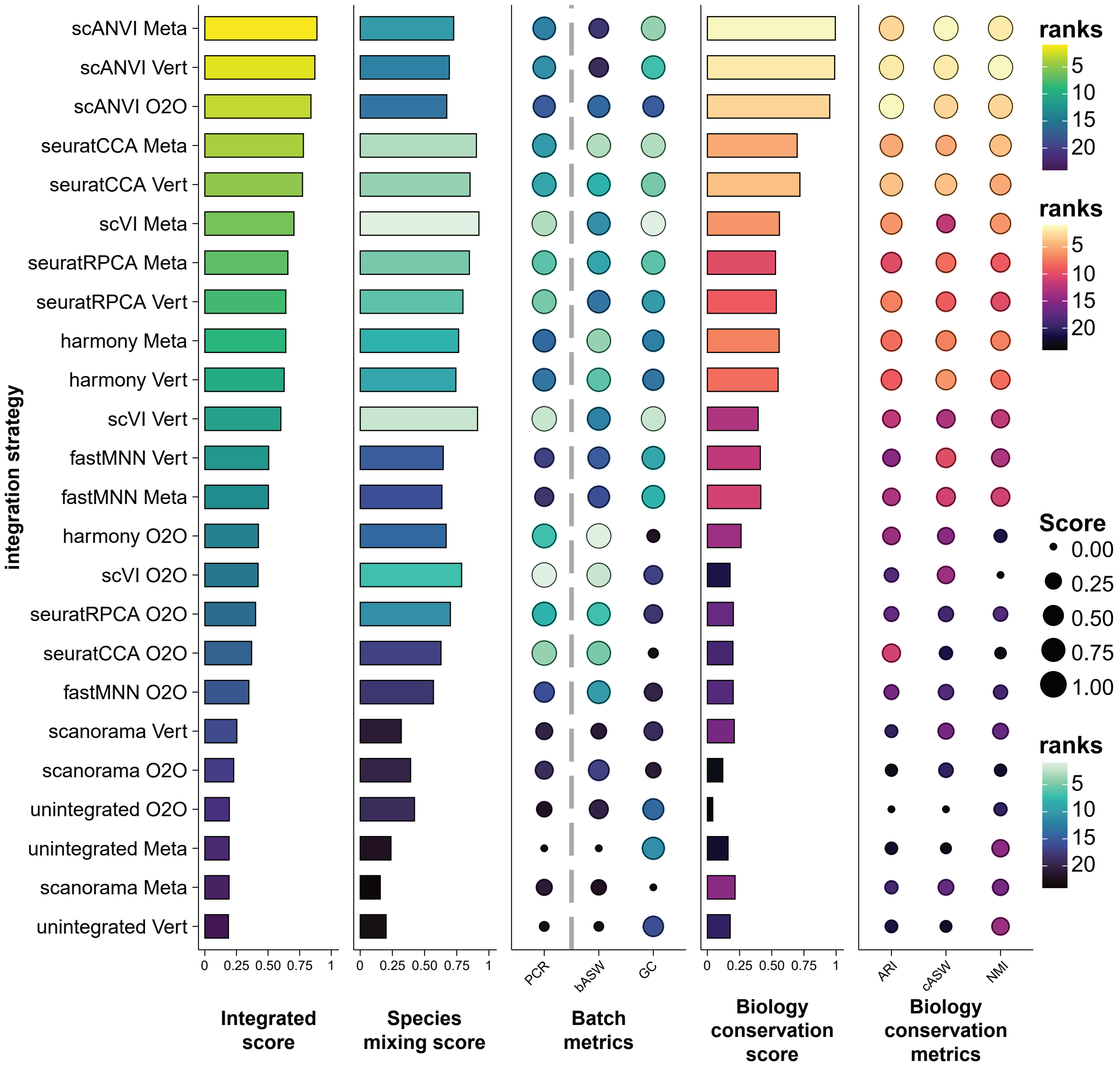

integration strategy
scANVI Meta
scANVI Vert
scANVI O2O
seuratCCA Meta
seuratCCA Vert
scVI Meta
seuratRPCA Meta
seuratRPCA Vert
harmony Meta
harmony Vert
scVI Vert
fastMNN Vert
fastMNN Meta
harmony O2O
scVI O2O
seuratRPCA O2O
seuratCCA O2O
fastMNN O2O
scanorama Vert
scanorama O2O
unintegrated O2O
unintegrated Meta
scanorama Meta
unintegrated Vert
0 0.25 0.50 0.75 1
Integrated score
Species mixing score
PCR
bASW
GC
Batch metrics
Biology conservation score
ARI
cASW
NMI
Biology conservation metrics
ranks
5
10
15
20
Score
0.00
0.25
0.50
0.75
1.00

**Figure S2: Vertebrate Pancreas Dataset Integration Comparison Scores**

Plots showing the breakdown of benchmarking scores for the integration of human, macaque, mouse, frog and zebrafish heart datasets. Integration method (tool+orthology method) shown on the left, ordered by overall integrated score. Metrics left to right: Integrated score, overall score averaging the species mixing and biological conservation scores; Species mixing score, an average of the metrics measuring the mixing of cells across species; Batch metrics, three metrics (PCR, bASW and GC) that denote if cells group by batch (in this case species); Biology conservation score, an average of metrics measuring the grouping of cells by their given cell type annotation; Biological conservation metrics, three metrics (ARI, cASW and NMI) that make up the Biological conservation score. The orthology methods are abbreviated as follows: O2O = 1:1 orthologs only; Vert = orthogroup recoding at the Vertebrata EggNOG level; Meta = orthogroup recoding at the Metazoa EggNOG level.

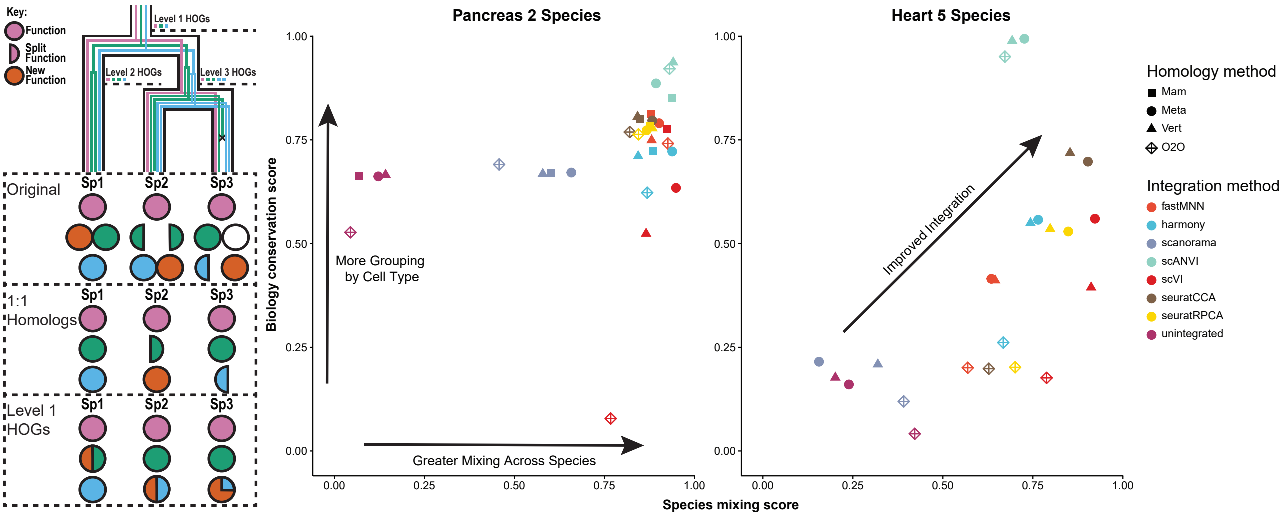

Key:
Function
Split Function
New Function
Level 1 HOGs
Level 2 HOGs
Level 3 HOGs
Original
1:1 Homologs
Level 1 HOGs
Sp1
Sp2
Sp3
Pancreas 2 Species
Heart 5 Species
Biology conservation score
Species mixing score
More Grouping by Cell Type
Greater Mixing Across Species
Improved Integration
Homology method
Mam
Meta
Vert
O2O
Integration method
fastMNN
harmony
scanorama
scANVI
scVI
seuratCCA
seuratRPCA
unintegrated

**Figure S3: Orthogroup Recoding Outperforms 1:1 Ortholog Integration**

A) Diagram illustrating the logic of orthogroup recoding, showing a species tree with gene relationships embedded within, the functions/expression patterns are symbolised with circles at the bottom. Three sets of circles indicate the preserved expressional information under the original data, 1:1 ortholog selection and orthogroup recoding. B) Scatter plots showing the integration scores calculated using scIB and BENGAL metrics to benchmark species integration methods. X-axis show the biological conservation score indicating how well cells grouped by cell type and the y-axis shows the species grouping score indicating how well the cells mixed across species. Scores are scaled 0-1 based on the best and worst performing methods. 7 integration methods are tested, represented by colours in the plots. 4 homology method are tested, represented by different shapes in the plots. I) benchmarking using H. sapiens and M. musculus pancreas data. Ii) benchmarking using H. sapiens, M. musculus, M. mulatta, X. tropicalis and D. rerio heart datasets.

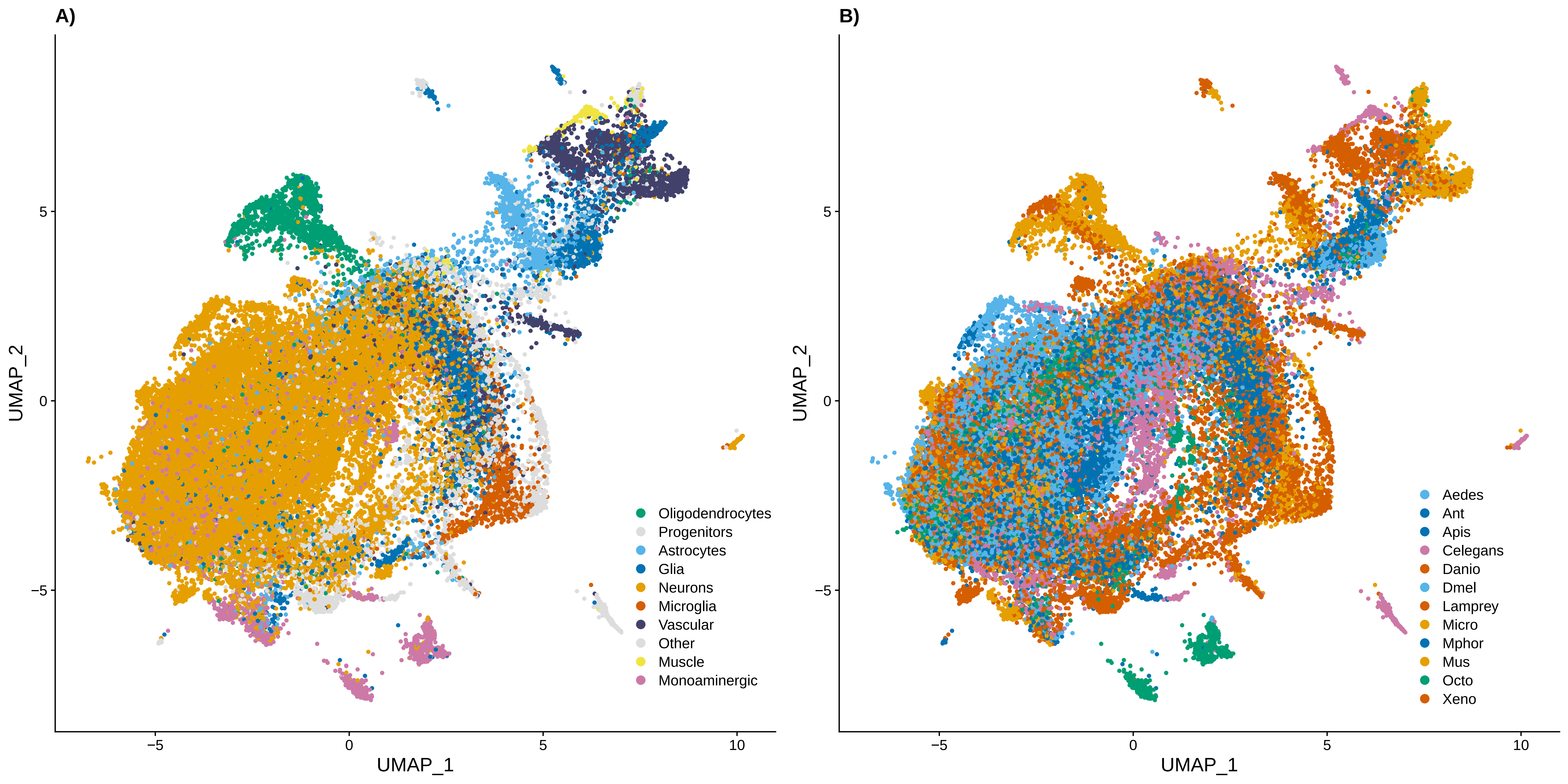
A)
B)
UMAP_1
UMAP_2
Oligodendrocytes
Progenitors
Astrocytes
Glia
Neurons
Microglia
Vascular
Other
Muscle
Monoaminergic
Aedes
Ant
Apis
Celegans
Danio
Dmel
Lamprey
Micro
Mphor
Mus
Octo
Xeno

**Figure S4: Bilaterian Brain Datasets Cluster by Broad Cell Type not by Species**

UMAPs of 12 bilaterian brain datasets integrated with orthogroup recoding using Seurat RPCA. A) Colour coded by broad cell type annotations. B) Colour coded by species.

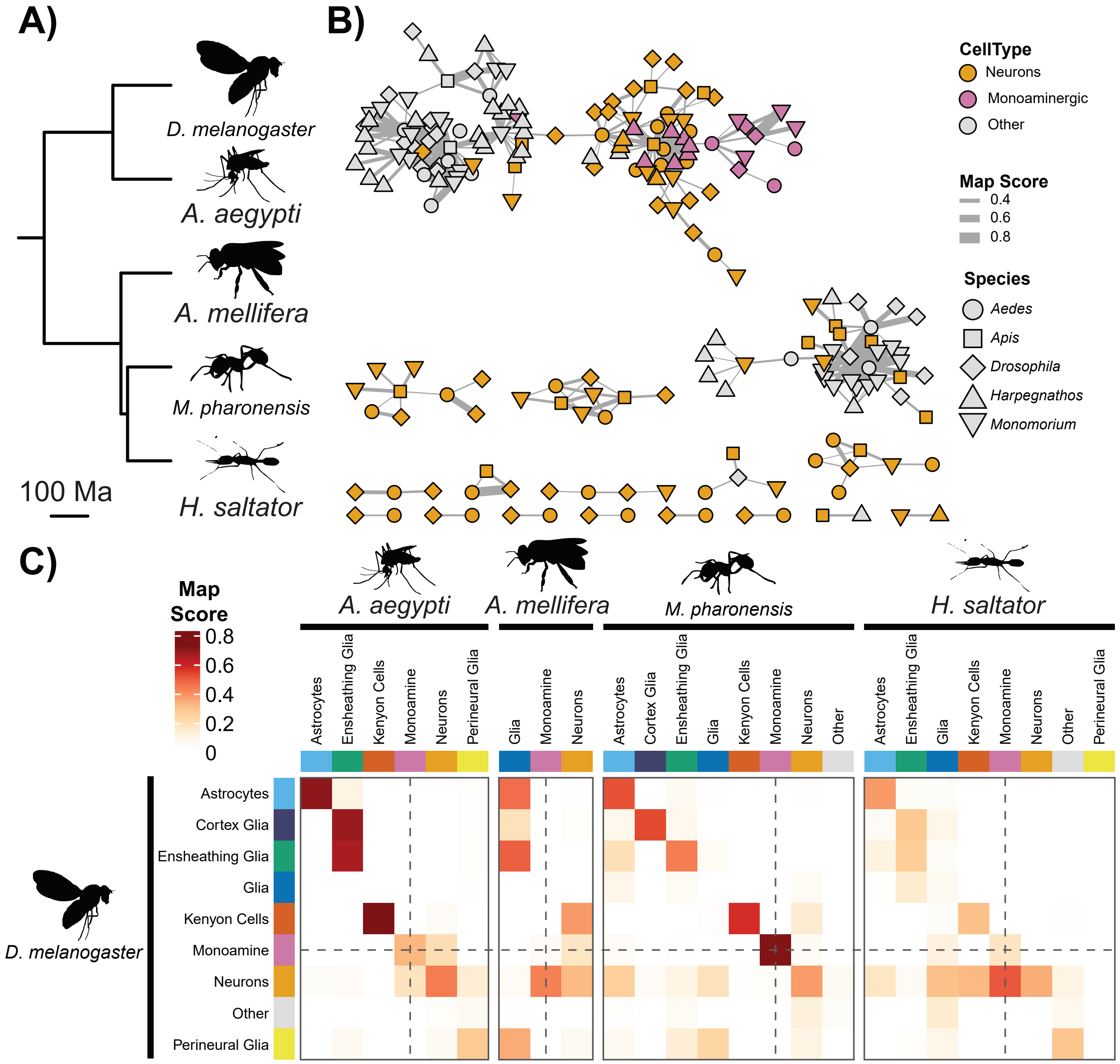

A)
B)
C)
D. melanogaster
A. aegypti
A. mellifera
M. pharonensis
H. saltator
100 Ma
CellType
Neurons
Monoaminergic
Other
Map Score
0.4
0.6
0.8
Species
Aedes
Apis
Drosophila
Harpegnathos
Monomorium
Map Score
0.8
0.6
0.4
0.2
0
Astrocytes
Ensheathing Glia
Kenyon Cells
Monoamine
Neurons
Perineural Glia
Glia
Cortex Glia
Other

**Figure S5: Insect Brain Monoaminergic Neurons Align with SAMap**

A) Time-scaled phylogeny of species used in the insect brain dataset. B) Network of SAMap mapping scores between individual cell clusters. Nodes represent cell clusters and edges represent SAMap mapping scores. Edge thickness denoted the mapping strength, scores below 0.2 are omitted. Nodes are colour coded: purple = monoaminergic; orange = neuron; white = other. C) Heatmap of SAMap mapping scores. Cell clusters have been summarised into broad groupings. Dashed lines follow the row/columns of monoaminergic groupings. Silhouettes from Phylopic.org: silhouette images are by Mattia Menchetti (*Apis mellifera*), Ramiro Morales-Hojas (*Drosophila americana* as *Drosophila melanogaster*), Richard J. Harris (*Aedes albopictus* as *Aedes aegpyti*, *Solenopsis invicta* as *Monomorium pharaonis*), and Vijay Karthick (*Harpegnathos saltator*).

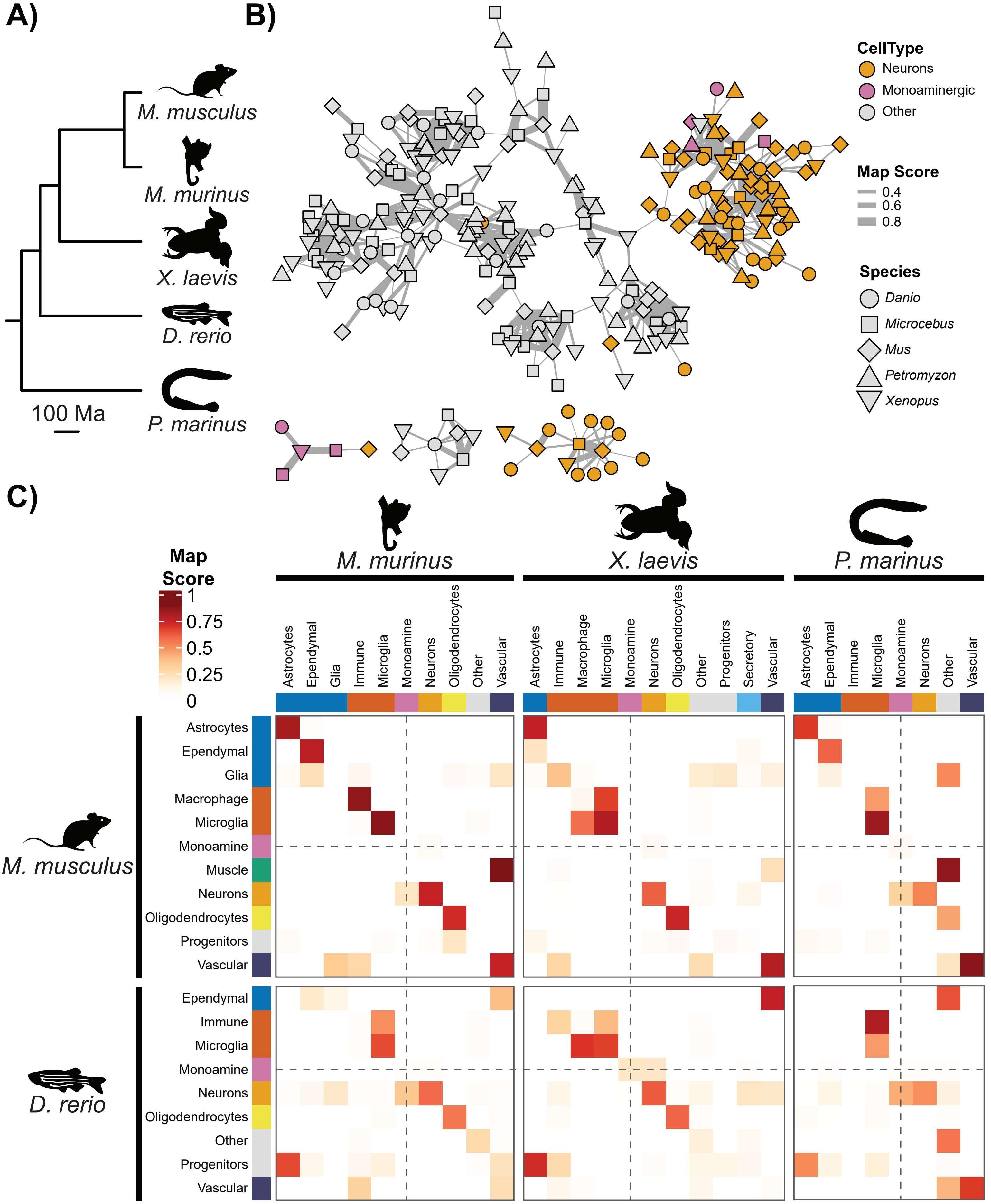

A)
B)
C)
M. musculus
M. murinus
X. laevis
D. rerio
P. marinus
100 Ma
CellType
Neurons
Monoaminergic
Other
Map Score
0.4
0.6
0.8
Species
Danio
Microcebus
Mus
Petromyzon
Xenopus
Map Score
1
0.75
0.5
0.25
0
Astrocytes
Ependymal
Glia
Immune
Microglia
Monoamine
Neurons
Oligodendrocytes
Other
Vascular
Macrophage
Progenitors
Secretory
Muscle

**Figure S6: Vertebrate Brain Monoaminergic Neurons do not Cluster with SAMap**

A) Time-scaled phylogeny of species used in the vertebrate brain dataset. B) Network of SAMap mapping scores between individual cell clusters. Nodes represent cell clusters and edges represent SAMap mapping scores. Edge thickness denoted the mapping strength, scores below 0.2 are omitted. Nodes are colour coded: purple = monoaminergic; orange = neuron; white = other. C) Heatmap of SAMap mapping scores. Cell clusters have been summarised into broad groupings. Dashed lines follow the row/columns of monoaminergic groupings. Silhouettes from Phylopic.org: silhouette images are by Daniel Jaron (*Mus musculus*), Jake Warner (*Danio rerio*), and others (*Microcebus murinus*, *Petromyzon marinus*, *Xenopus laevis*).

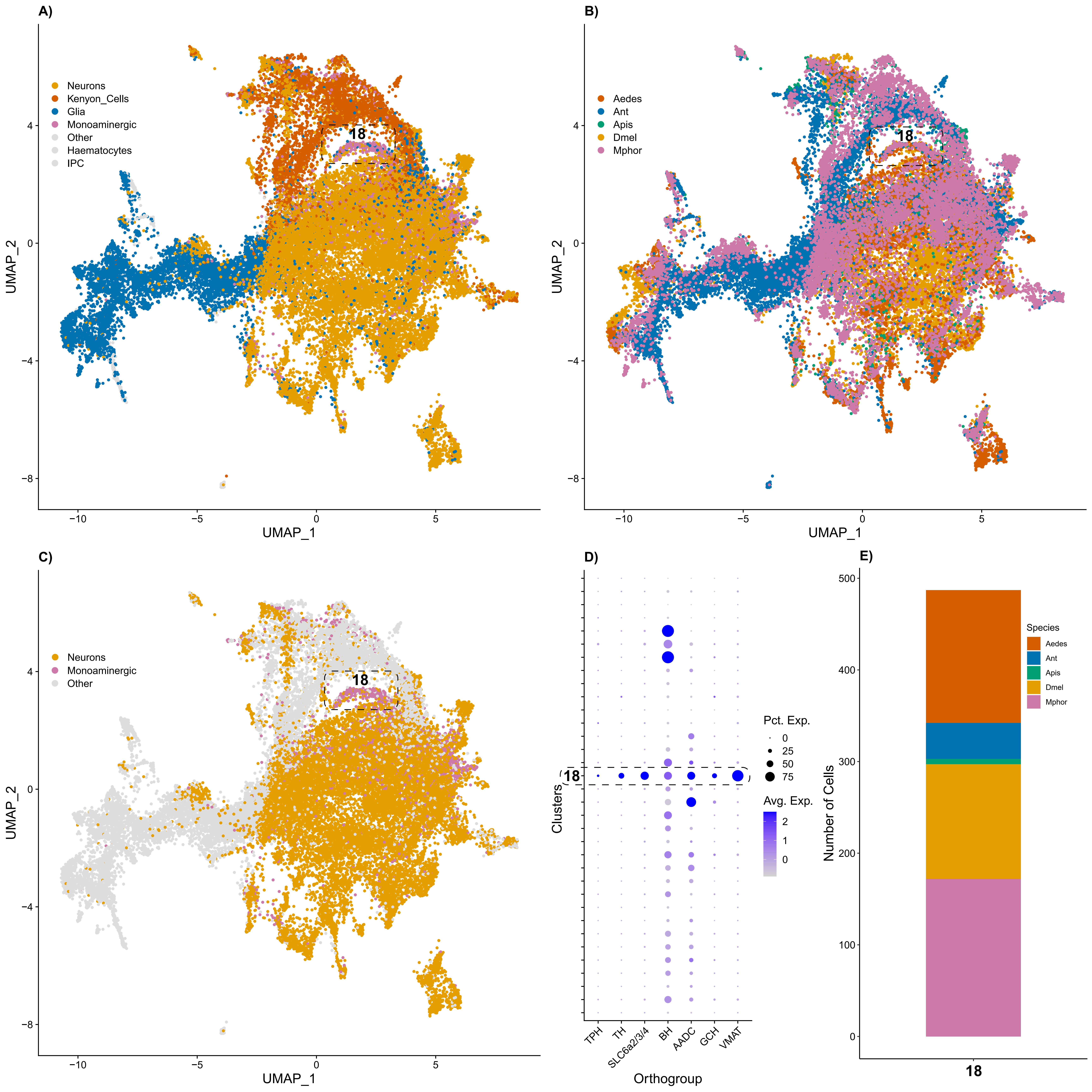
A)
Neurons
Kenyon_Cells
Glia
Monoaminergic
Other
Haematocytes
IPC
18
UMAP_1
UMAP_2
B)
Aedes
Ant
Apis
Dmel
Mphor
18
UMAP_1
UMAP_2
C)
Neurons
Monoaminergic
Other
18
UMAP_1
UMAP_2
D)
18
Clusters
Pct. Exp.
0
25
50
75
Avg. Exp.
2
1
0
TPH
TH
SLC6a2/3/4
BH
AADC
GCH
VMAT
Orthogroup
E)
Species
Aedes
Ant
Apis
Dmel
Mphor
Number of Cells
18

**Figure S7: Insect Brain Monoaminergic Neurons Cluster Across Species**

Plots detailing the clustering and expression of monoaminergic cells from 5 insect brain datasets under Orthogroup Recoding. A) UMAP colour coded by broad cell type annotations. B) UMAP colour coded by species. C) RPCA integrated UMAP, colour coded by annotation. Monoaminergic clusters highlighted with dashed boxes and labelled. D) Dot plot showing the expression of monoaminergic orthogroups across clusters. E) Bar plot of the species makeup of monoaminergic clusters. Gene expression was recoded using the Metazoa level of EggNOG annotations.

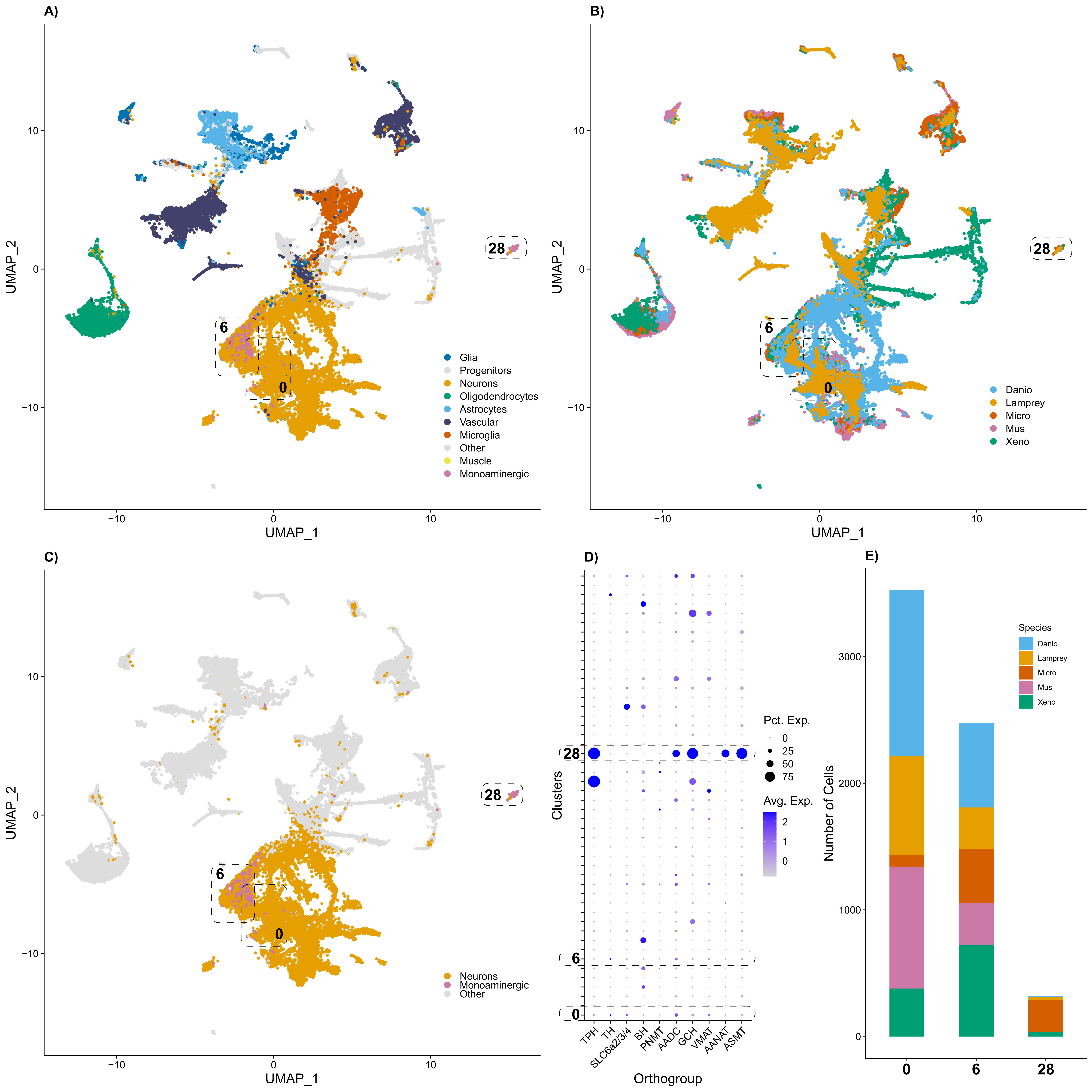

A)
B)
C)
D)
E)
UMAP_1
UMAP_2
Glia
Progenitors
Neurons
Oligodendrocytes
Astrocytes
Vascular
Microglia
Other
Muscle
Monoaminergic
Danio
Lamprey
Micro
Mus
Xeno
Neurons
Monoaminergic
Other
Pct. Exp.
Avg. Exp.
Clusters
Orthogroup
TPH
TH
SLC6a2/3/4
BH
PNMT
AADC
GCH
VMAT
AANAT
ASMT
Species
Number of Cells
0
6
28

**Figure S8: Vertebrate Brain Monoaminergic Neurons show heterogenous clustering**

Plots detailing the clustering and expression of monoaminergic cells from 5 vertebrate brain datasets under Orthogroup Recoding. A) UMAP colour coded by broad cell type annotations. B) UMAP colour coded by species. C) RPCA integrated UMAP, colour coded by annotation. Monoaminergic clusters highlighted with dashed boxes and labelled. D) Dot plot showing the expression of monoaminergic orthogroups across clusters. E) Bar plot of the species makeup of monoaminergic clusters. Gene expression was recoded using the Metazoa level of EggNOG annotations.

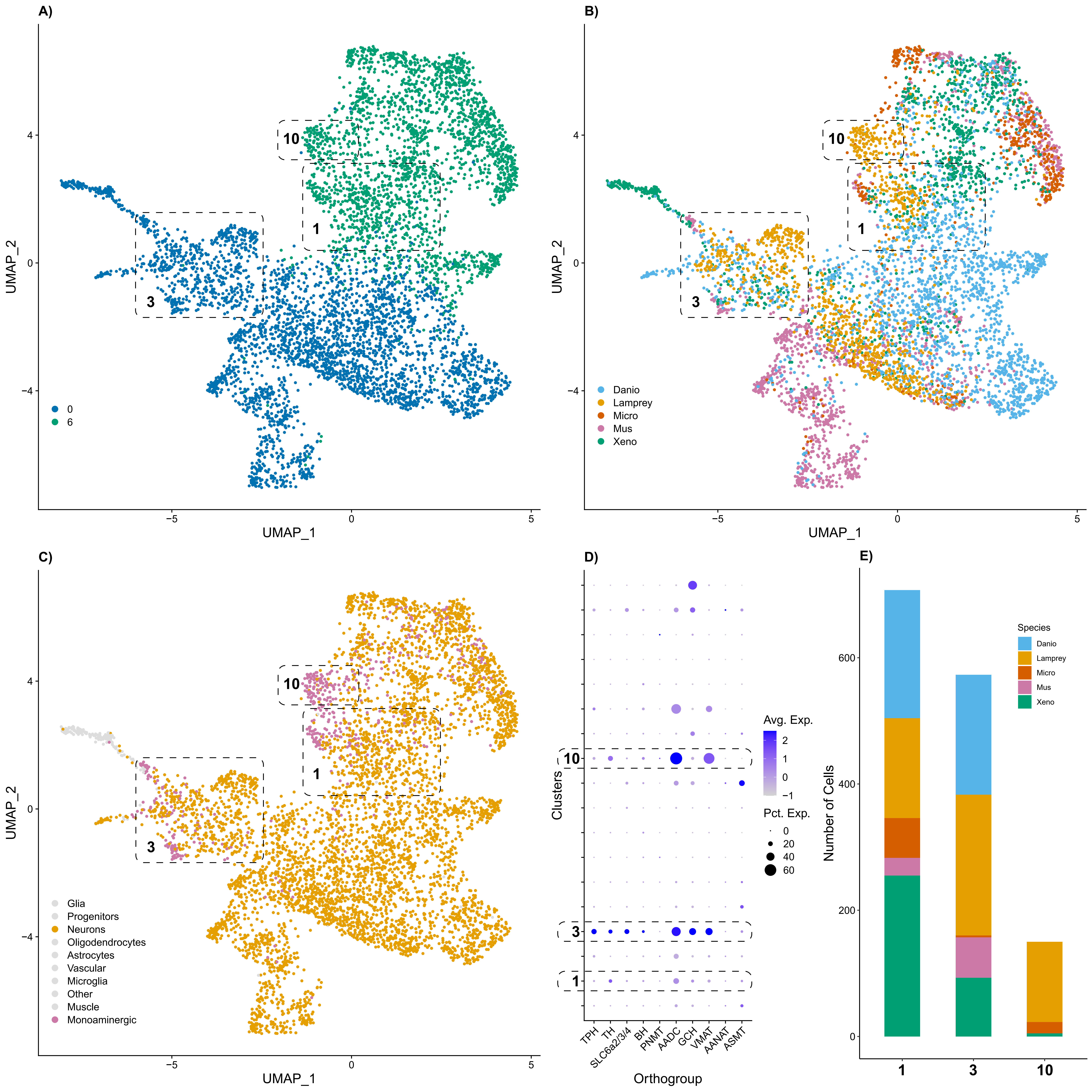

A)
B)
C)
D)
E)
UMAP_1
UMAP_2
10
1
3
0
6
Danio
Lamprey
Micro
Mus
Xeno
Glia
Progenitors
Neurons
Oligodendrocytes
Astrocytes
Vascular
Microglia
Other
Muscle
Monoaminergic
Clusters
Orthogroup
TPH
TH
SLC6a2/3/4
BH
PNMT
AADC
GCH
VMAT
AANAT
ASMT
Avg. Exp.
Pct. Exp.
Species
Number of Cells

**Figure S9: Vertebrate Brain Monoaminergic Neurons form Subclusters within Neurons**

A) Plots detailing the sub-clustering and expression of monoaminergic cells from 5 vertebrate brain datasets under Orthogroup Recoding. A) UMAP colour coded by main clusters used for the sub-clustering. B) UMAP colour coded by species. C) RPCA integrated UMAP, colour coded by annotation. Monoaminergic subclusters highlighted with dashed boxes and labelled. D) Dot plot showing the expression of monoaminergic orthogroups across subclusters. E) Bar plot of the species makeup of monoaminergic subclusters. Gene expression was recoded using the Metazoa level of EggNOG annotations.

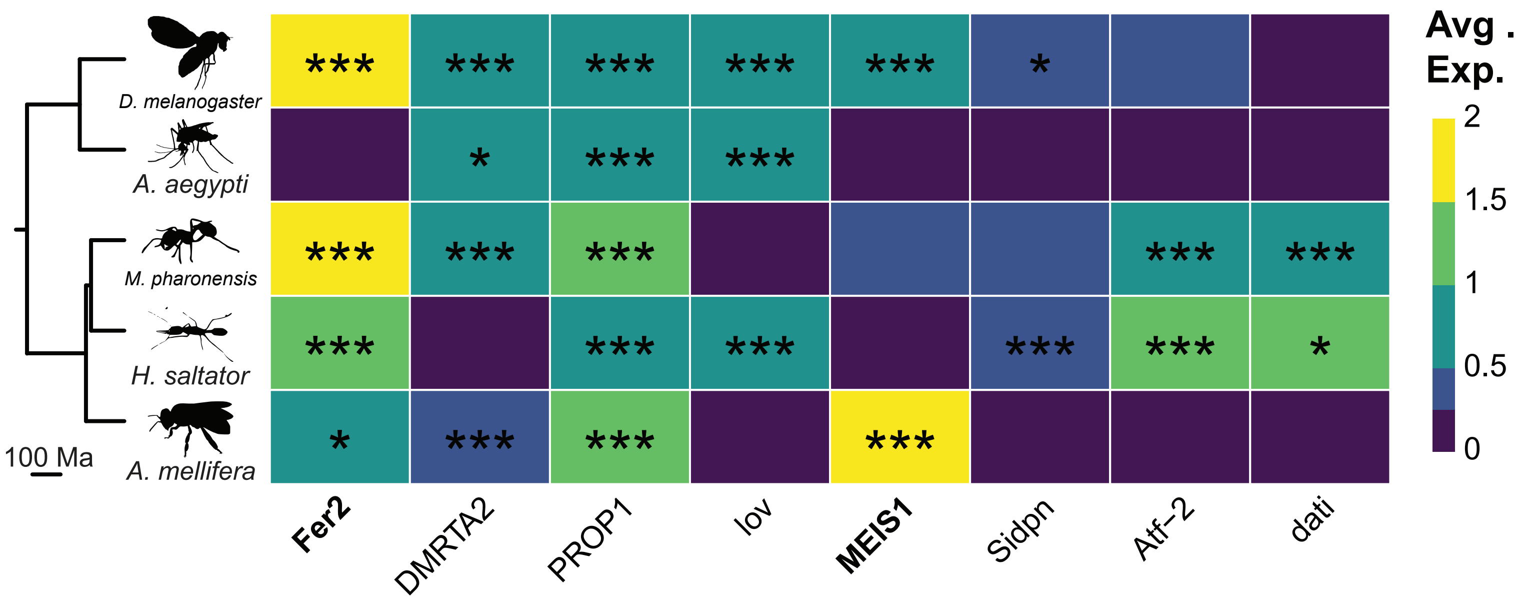
D. melanogaster
A. aegypti
M. pharonensis
H. saltator
A. mellifera
100 Ma
Fer2
DMRTA2
PROP1
lov
MEIS1
Sidpn
Atf-2
dati
Avg . Exp.
2
1.5
1
0.5
0

**Figure S10: Insect Brain Monoaminergic Neurons share a core transcription factor profile**

Heatmaps of differentially expressed transcription factor orthogroups shared across species in the monoaminergic cluster from the insect brain integration. Species tree and icons displayed on the left indicating the species included. Orthogroup names shown along the bottom, orthogroups listed in table 1 are in bold. Orthogroups are included when they are significantly expressed in two or more species. The colour code corresponds to log fold change expression < 0.25 lfc is omitted and * indicates p.adj<0.05. Silhouettes from Phylopic.org: silhouette images are by Mattia Menchetti (*Apis mellifera*), Ramiro Morales-Hojas (*Drosophila americana* as *Drosophila melanogaster*), Richard J. Harris (*Aedes albopictus* as *Aedes aegpyti*, *Solenopsis invicta* as *Monomorium pharaonis*), and Vijay Karthick (*Harpegnathos saltator*).

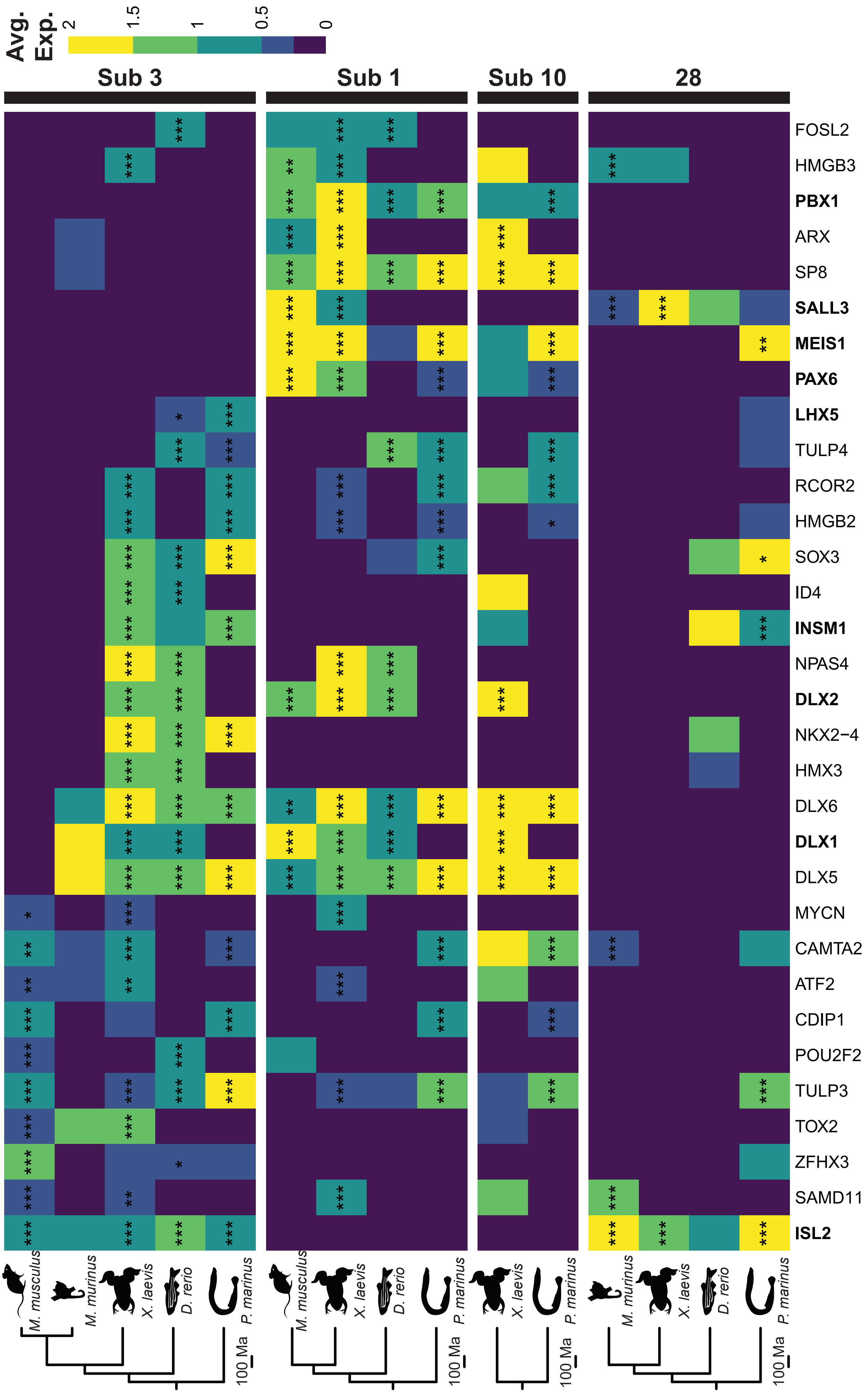

Avg. Exp.
2
1.5
1
0.5
0
Sub 3
Sub 1
Sub 10
28
FOSL2
HMGB3
PBX1
ARX
SP8
SALL3
MEIS1
PAX6
LHX5
TULP4
RCOR2
HMGB2
SOX3
ID4
INSM1
NPAS4
DLX2
NKX2−4
HMX3
DLX6
DLX1
DLX5
MYCN
CAMTA2
ATF2
CDIP1
POU2F2
TULP3
TOX2
ZFHX3
SAMD11
ISL2
M. musculus
M. murinus
X. laevis
D. rerio
P. marinus
100 Ma

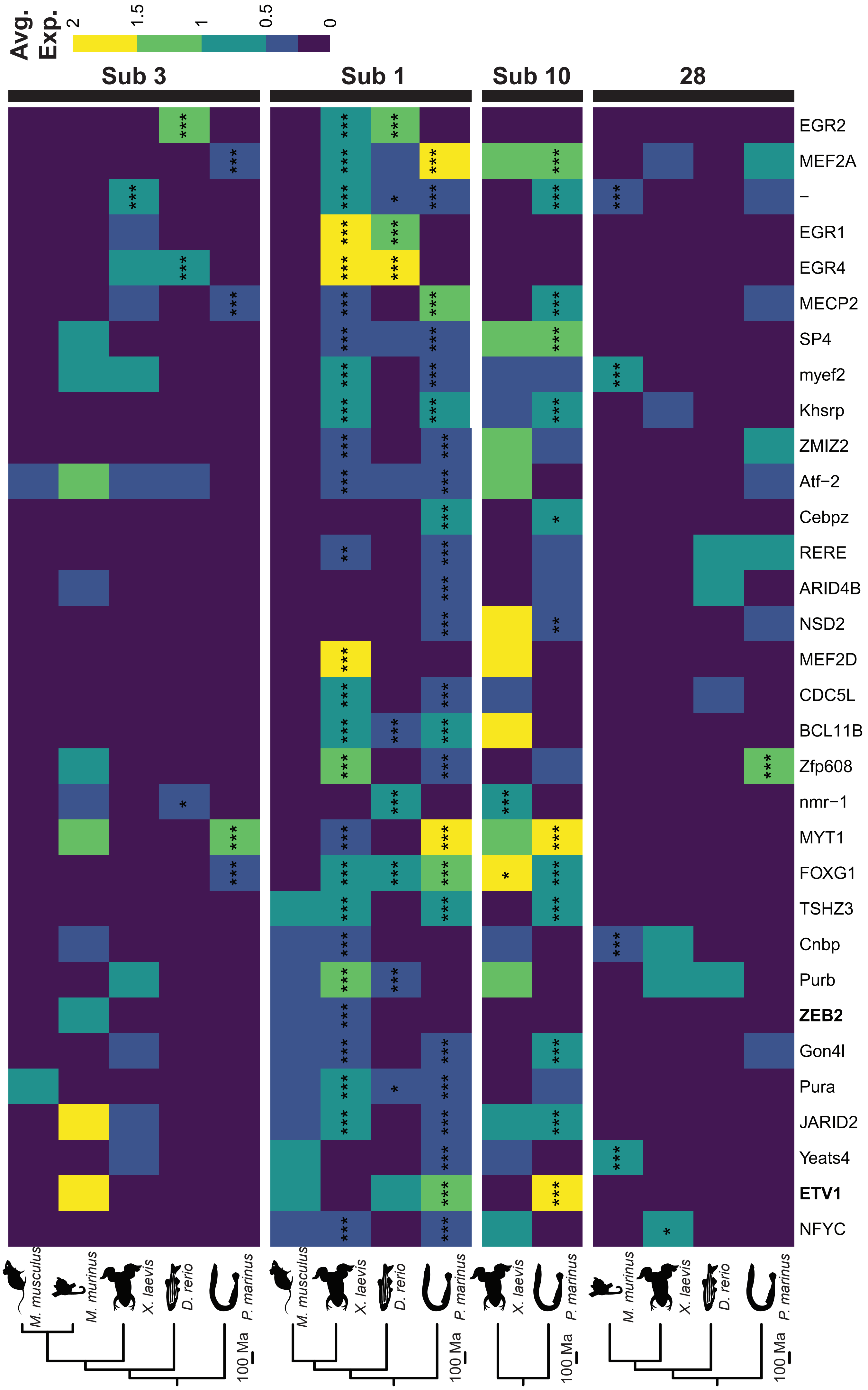

Avg. Exp.
2
1.5
1
0.5
0
Sub 3
Sub 1
Sub 10
28
EGR2
MEF2A
–
EGR1
EGR4
MECP2
SP4
myef2
Khsrp
ZMIZ2
Atf−2
Cebpz
RERE
ARID4B
NSD2
MEF2D
CDC5L
BCL11B
Zfp608
nmr−1
MYT1
FOXG1
TSHZ3
Cnbp
Purb
ZEB2
Gon4l
Pura
JARID2
Yeats4
ETV1
NFYC
M. musculus
M. murinus
X. laevis
D. rerio
P. marinus
100 Ma

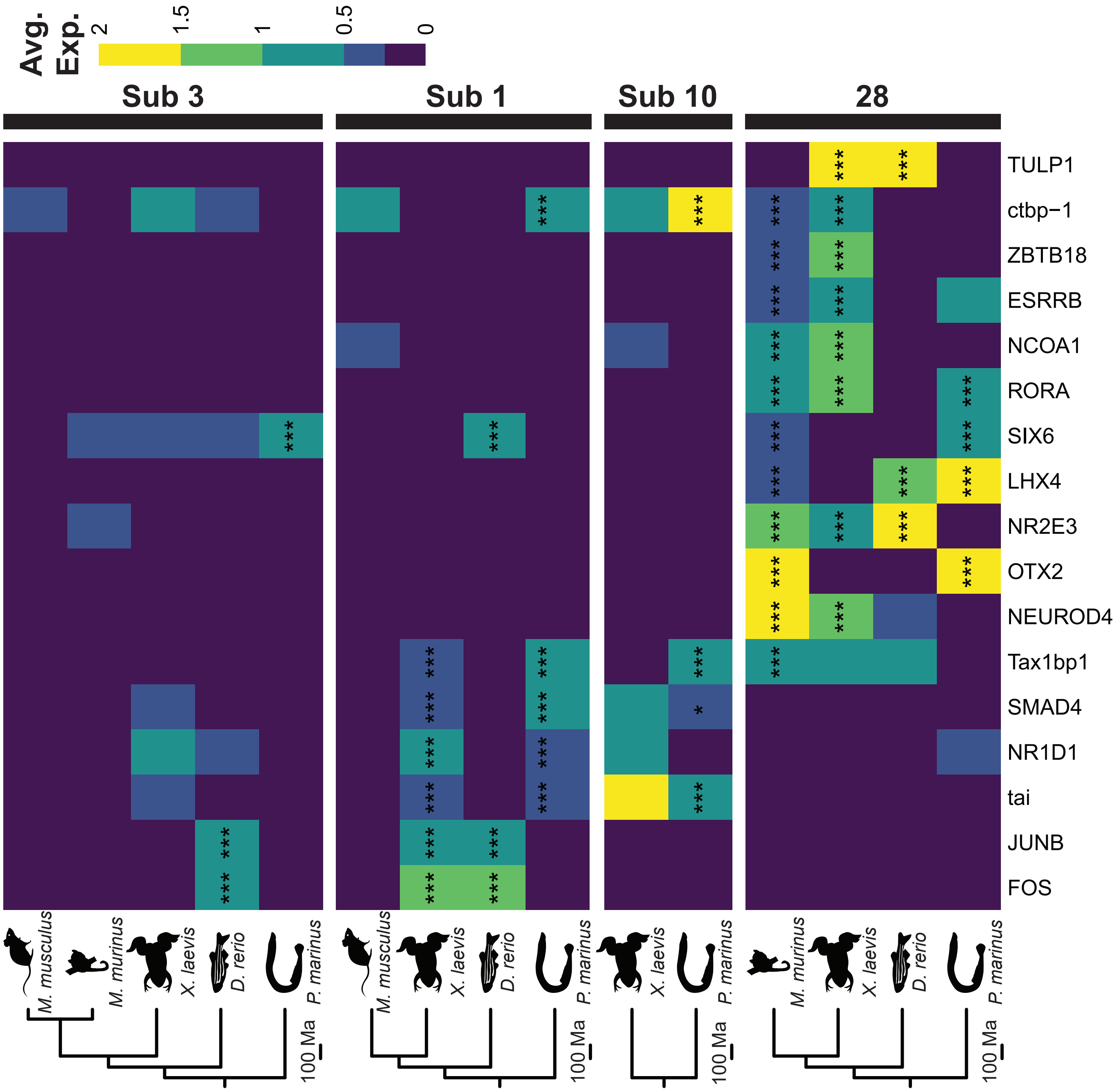

Avg. Exp.
2
1.5
1
0.5
0
Sub 3
Sub 1
Sub 10
28
TULP1
ctbp−1
ZBTB18
ESRRB
NCOA1
RORA
SIX6
LHX4
NR2E3
OTX2
NEUROD4
Tax1bp1
SMAD4
NR1D1
tai
JUNB
FOS
M. musculus
M. murinus
X. laevis
D. rerio
P. marinus
100 Ma

**Figure S11: Vertebrate Brain Monoaminergic Neurons do share some transcription factor expression**

Heatmaps of differentially expressed transcription factor orthogroups shared across species in the four monoaminergic clusters/subclusters of the vertebrate brain integration. Species trees and icons displayed on the left indicating the species included. Orthogroup names shown along the bottom, orthogroups listed in table 1 are in bold. Orthogroups are included when they are expressed in two or more species in a cluster/subcluster of interest. Orthogroups are mapped across all four monoaminergic clusters if shared in one cluster. The colour code corresponds to log fold change expression < 0.25 lfc is omitted and * indicates p.adj<0.05. Silhouettes from Phylopic.org: silhouette images are by Daniel Jaron (*Mus musculus*), Jake Warner (*Danio rerio*), and others (*Microcebus murinus*, *Petromyzon marinus*, *Xenopus laevis*).

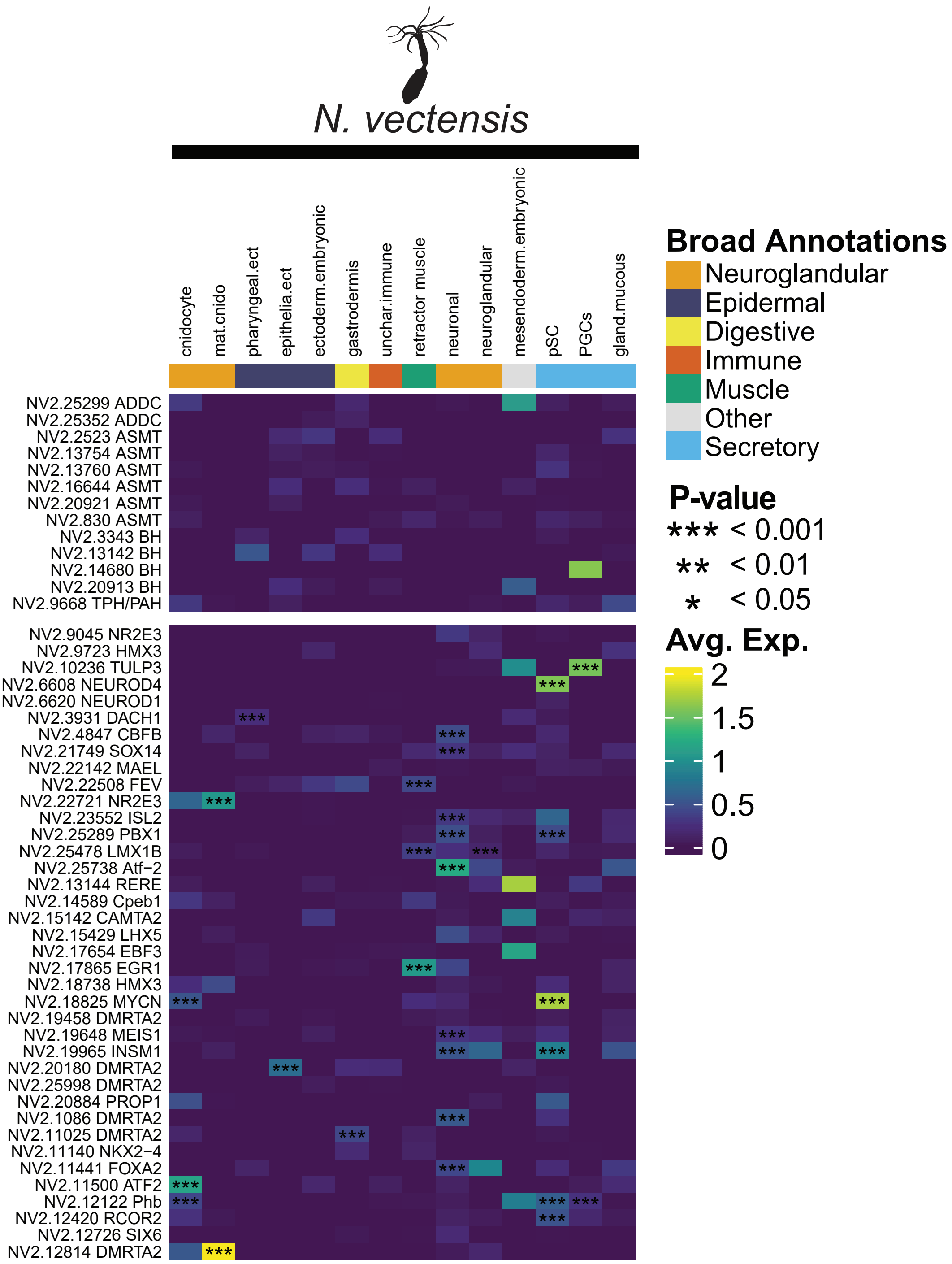

N. vectensis
cnidocyte
mat.cnido
pharyngeal.ect
epithelia.ect
ectoderm.embryonic
gastrodermis
unchar.immune
retractor muscle
neuronal
neuroglandular
mesendoderm.embryonic
pSC
PGCs
gland.mucous
NV2.25299 ADDC
NV2.25352 ADDC
NV2.2523 ASMT
NV2.13754 ASMT
NV2.13760 ASMT
NV2.16644 ASMT
NV2.20921 ASMT
NV2.830 ASMT
NV2.3343 BH
NV2.13142 BH
NV2.14680 BH
NV2.20913 BH
NV2.9668 TPH/PAH
NV2.9045 NR2E3
NV2.9723 HMX3
NV2.10236 TULP3
NV2.6608 NEUROD4
NV2.6620 NEUROD1
NV2.3931 DACH1
NV2.4847 CBFB
NV2.21749 SOX14
NV2.22142 MAEL
NV2.22508 FEV
NV2.22721 NR2E3
NV2.23552 ISL2
NV2.25289 PBX1
NV2.25478 LMX1B
NV2.25738 Atf−2
NV2.13144 RERE
NV2.14589 Cpeb1
NV2.15142 CAMTA2
NV2.15429 LHX5
NV2.17654 EBF3
NV2.17865 EGR1
NV2.18738 HMX3
NV2.18825 MYCN
NV2.19458 DMRTA2
NV2.19648 MEIS1
NV2.19965 INSM1
NV2.20180 DMRTA2
NV2.25998 DMRTA2
NV2.20884 PROP1
NV2.1086 DMRTA2
NV2.11025 DMRTA2
NV2.11140 NKX2−4
NV2.11441 FOXA2
NV2.11500 ATF2
NV2.12122 Phb
NV2.12420 RCOR2
NV2.12726 SIX6
NV2.12814 DMRTA2
Broad Annotations
Neuroglandular
Epidermal
Digestive
Immune
Muscle
Other
Secretory
P-value
*** < 0.001
** < 0.01
* < 0.05
Avg. Exp.
2
1.5
1
0.5
0

**Figure S12: *Nematostella vectensis* shows no co-ordinated expression of monoaminergic pathway genes or transcription factors**

Heatmap of monoaminergic pathway and transcription factor gene expression in *Nematostella vectensis*. Rows named by gene name plus orthogroup name. Colour coded by log fold change of expression. Significance of differential expression is indicated by *. Cell type annotations colour coded by broad grouping. Silhouettes from Phylopic.org: silhouette image by Jake Warner (Nematostella vectensis).

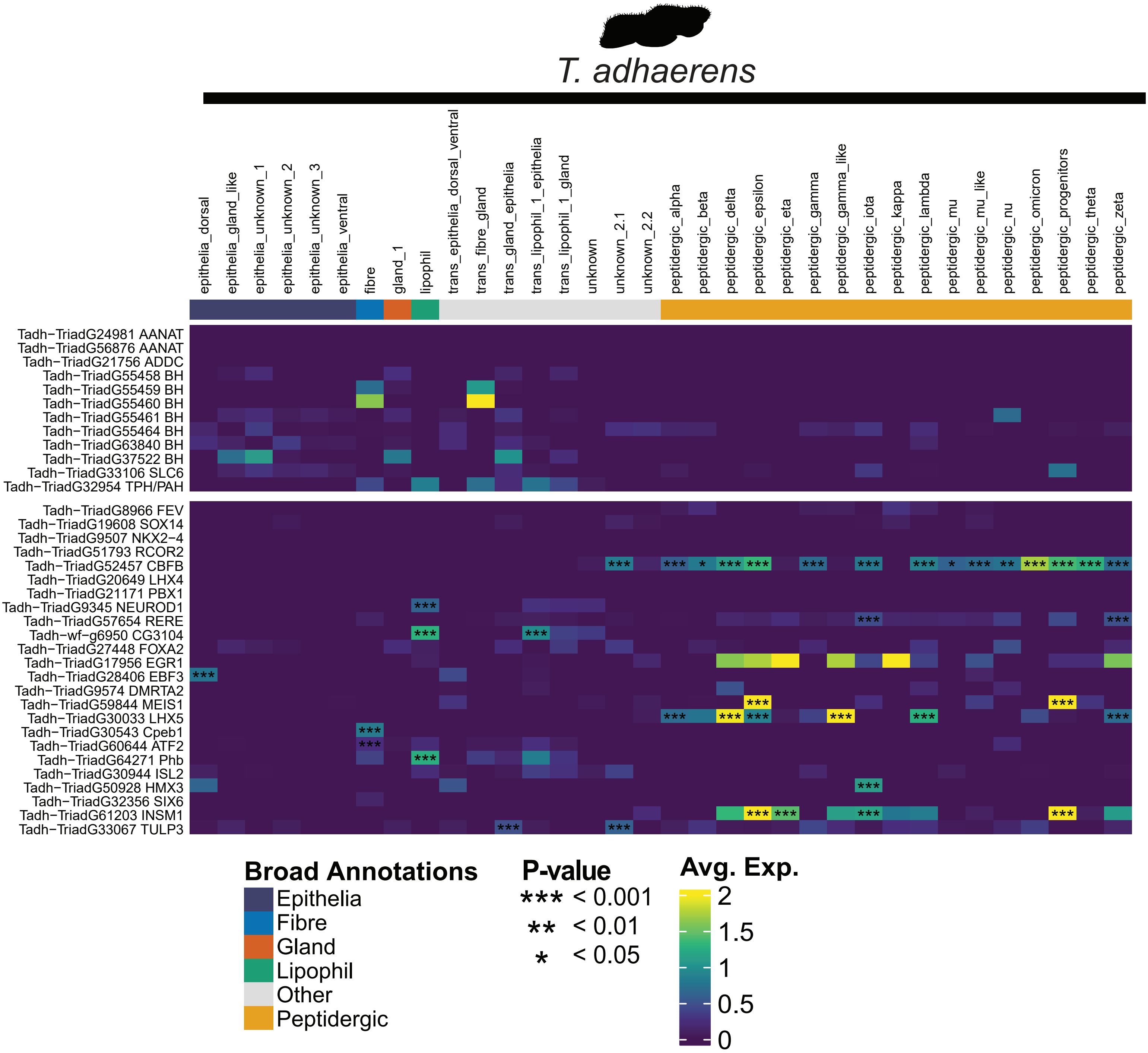
T. adhaerens
epithelia_dorsal
epithelia_gland_like
epithelia_unknown_1
epithelia_unknown_2
epithelia_unknown_3
epithelia_ventral
fibre
gland_1
lipophil
trans_epithelia_dorsal_ventral
trans_fibre_gland
trans_gland_epithelia
trans_lipophil_1_epithelia
trans_lipophil_1_gland
unknown
unknown_2.1
unknown_2.2
peptidergic_alpha
peptidergic_beta
peptidergic_delta
peptidergic_epsilon
peptidergic_eta
peptidergic_gamma
peptidergic_gamma_like
peptidergic_iota
peptidergic_kappa
peptidergic_lambda
peptidergic_mu
peptidergic_mu_like
peptidergic_nu
peptidergic_omicron
peptidergic_progenitors
peptidergic_theta
peptidergic_zeta
Tadh−TriadG24981 AANAT
Tadh−TriadG56876 AANAT
Tadh−TriadG21756 ADDC
Tadh−TriadG55458 BH
Tadh−TriadG55459 BH
Tadh−TriadG55460 BH
Tadh−TriadG55461 BH
Tadh−TriadG55464 BH
Tadh−TriadG63840 BH
Tadh−TriadG37522 BH
Tadh−TriadG33106 SLC6
Tadh−TriadG32954 TPH/PAH
Tadh−TriadG8966 FEV
Tadh−TriadG19608 SOX14
Tadh−TriadG9507 NKX2−4
Tadh−TriadG51793 RCOR2
Tadh−TriadG52457 CBFB
Tadh−TriadG20649 LHX4
Tadh−TriadG21171 PBX1
Tadh−TriadG9345 NEUROD1
Tadh−TriadG57654 RERE
Tadh−wf−g6950 CG3104
Tadh−TriadG27448 FOXA2
Tadh−TriadG17956 EGR1
Tadh−TriadG28406 EBF3
Tadh−TriadG9574 DMRTA2
Tadh−TriadG59844 MEIS1
Tadh−TriadG30033 LHX5
Tadh−TriadG30543 Cpeb1
Tadh−TriadG60644 ATF2
Tadh−TriadG64271 Phb
Tadh−TriadG30944 ISL2
Tadh−TriadG50928 HMX3
Tadh−TriadG32356 SIX6
Tadh−TriadG61203 INSM1
Tadh−TriadG33067 TULP3
Broad Annotations
Epithelia
Fibre
Gland
Lipophil
Other
Peptidergic
P-value
*** < 0.001
** < 0.01
* < 0.05
Avg. Exp.
2
1.5
1
0.5
0

**Figure S13: *Trichoplax adhearens* shows no co-ordinated expression of monoaminergic pathway genes or transcription factors**

Heatmap of monoaminergic pathway and transcription factor gene expression in *Trichoplax adhearens*. Rows named by gene name plus orthogroup name. Colour coded by log fold change of expression. Significance of differential expression is indicated by *. Cell type annotations colour coded by broad grouping. Silhouettes from Phylopic.org: silhouette image by Yan Wong (Trichoplax adhaerens).

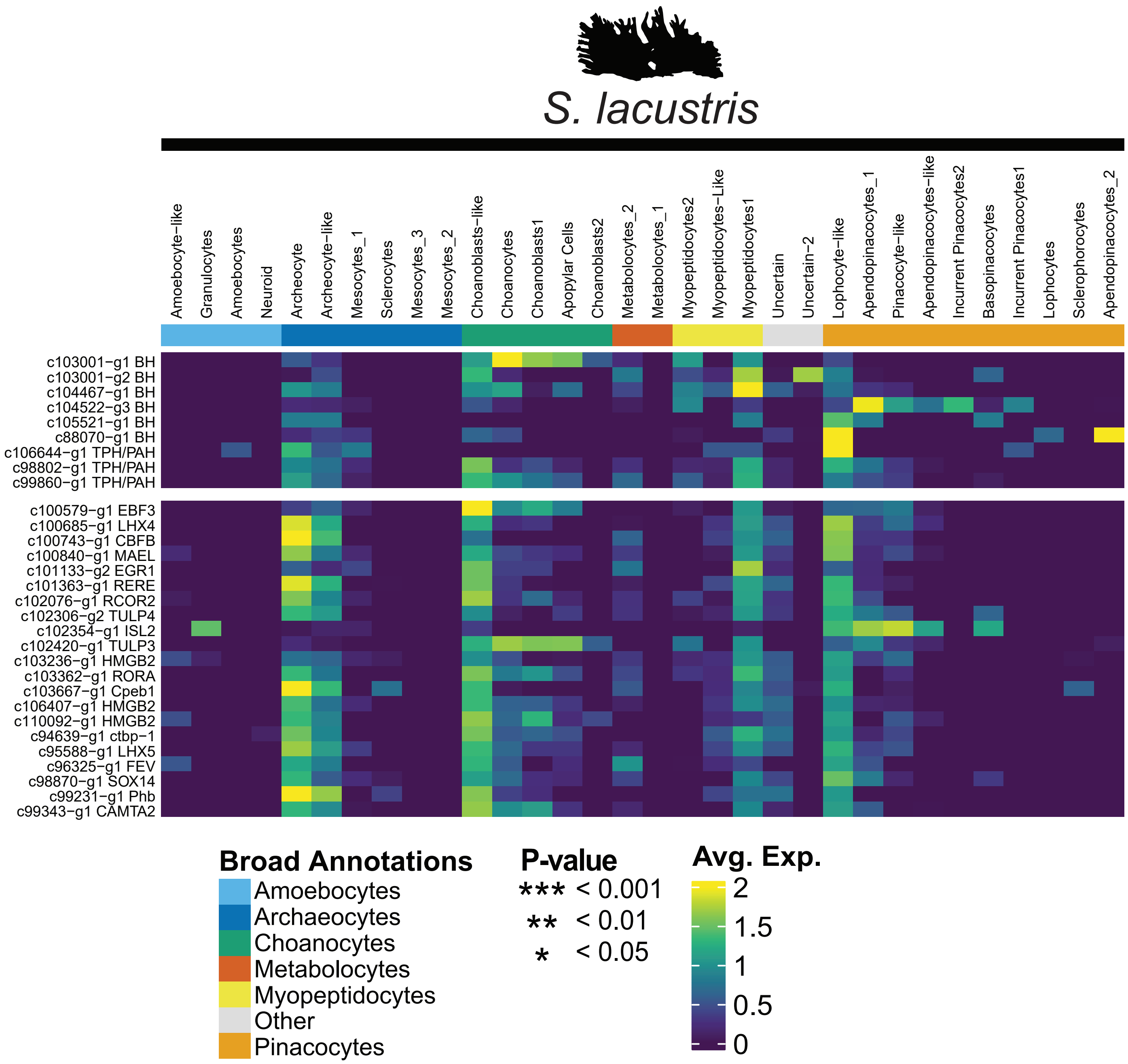
S. lacustris
Amoebocyte−like
Granulocytes
Amoebocytes
Neuroid
Archeocyte
Archeocyte−like
Mesocytes_1
Sclerocytes
Mesocytes_3
Mesocytes_2
Choanoblasts−like
Choanocytes
Choanoblasts1
Apopylar Cells
Choanoblasts2
Metabolocytes_2
Metabolocytes_1
Myopeptidocytes2
Myopeptidocytes−Like
Myopeptidocytes1
Uncertain
Uncertain−2
Lophocyte−like
Apendopinacocytes_1
Pinacocyte−like
Apendopinacocytes−like
Incurrent Pinacocytes2
Basopinacocytes
Incurrent Pinacocytes1
Lophocytes
Sclerophorocytes
Apendopinacocytes_2
c103001−g1 BH
c103001−g2 BH
c104467−g1 BH
c104522−g3 BH
c105521−g1 BH
c88070−g1 BH
c106644−g1 TPH/PAH
c98802−g1 TPH/PAH
c99860−g1 TPH/PAH
c100579−g1 EBF3
c100685−g1 LHX4
c100743−g1 CBFB
c100840−g1 MAEL
c101133−g2 EGR1
c101363−g1 RERE
c102076−g1 RCOR2
c102306−g2 TULP4
c102354−g1 ISL2
c102420−g1 TULP3
c103236−g1 HMGB2
c103362−g1 RORA
c103667−g1 Cpeb1
c106407−g1 HMGB2
c110092−g1 HMGB2
c94639−g1 ctbp−1
c95588−g1 LHX5
c96325−g1 FEV
c98870−g1 SOX14
c99231−g1 Phb
c99343−g1 CAMTA2
Broad Annotations
Amoebocytes
Archaeocytes
Choanocytes
Metabolocytes
Myopeptidocytes
Other
Pinacocytes
P-value
*** < 0.001
** < 0.01
* < 0.05
Avg. Exp.
2
1.5
1
0.5
0

**Figure S14: *Spongilla lacustris* shows no co-ordinated expression of monoaminergic pathway genes or transcription factors**

Heatmap of monoaminergic pathway and transcription factor gene expression in *Spongilla lacustris*. Rows named by gene name plus orthogroup name. Colour coded by log fold change of expression. Significance of differential expression is indicated by *. Cell type annotations colour coded by broad grouping. Silhouettes from Phylopic.org: silhouette image *Isodictya grandis* as *Spongilla lacustris*.

A)

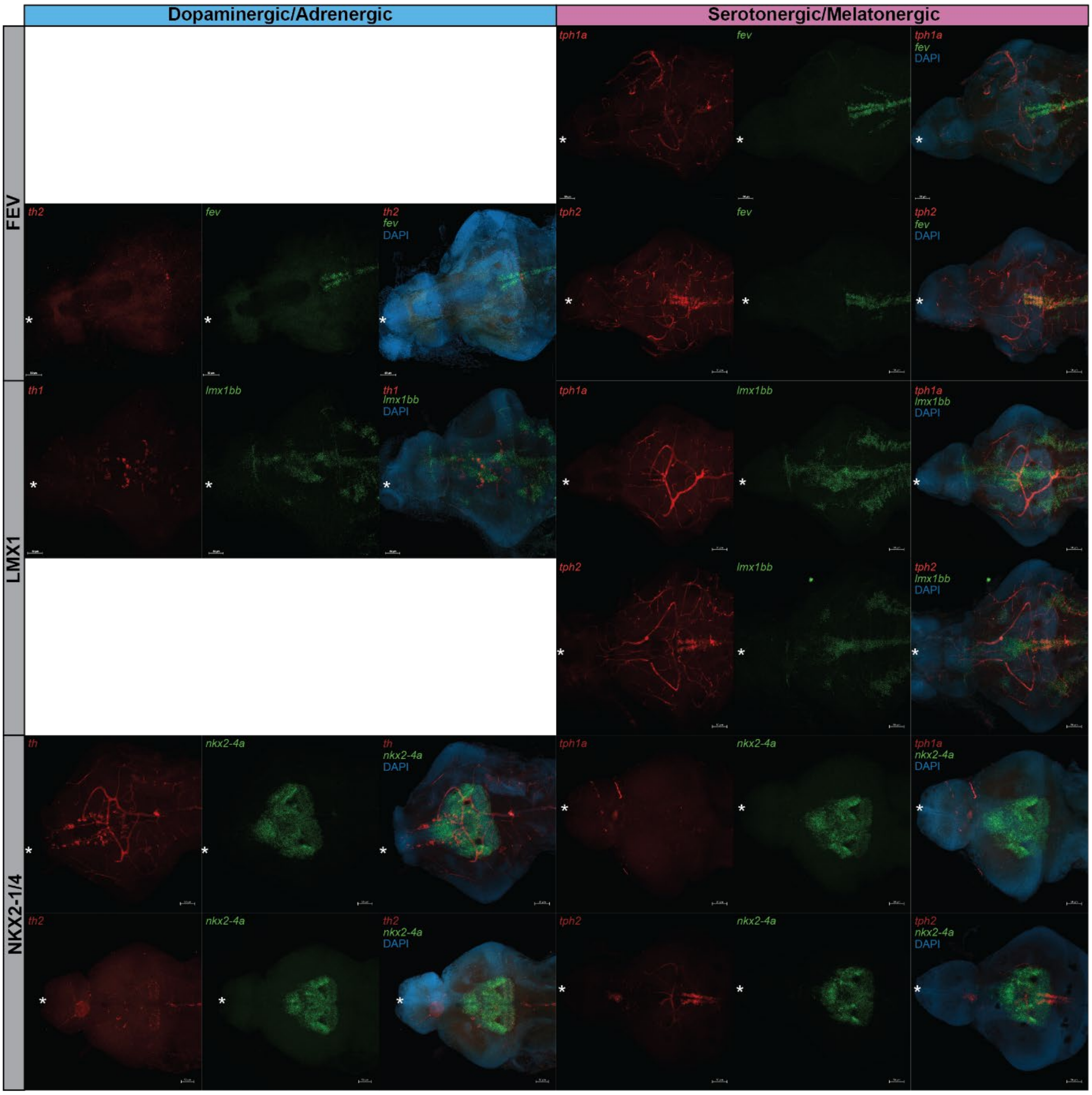

Dopaminergic/Adrenergic
Serotonergic/Melatonergic
FEV
LMX1
NKX2-1/4
tph1a
fev
DAPI
th2
tph2
th1
lmx1bb
th
nkx2-4a

B)

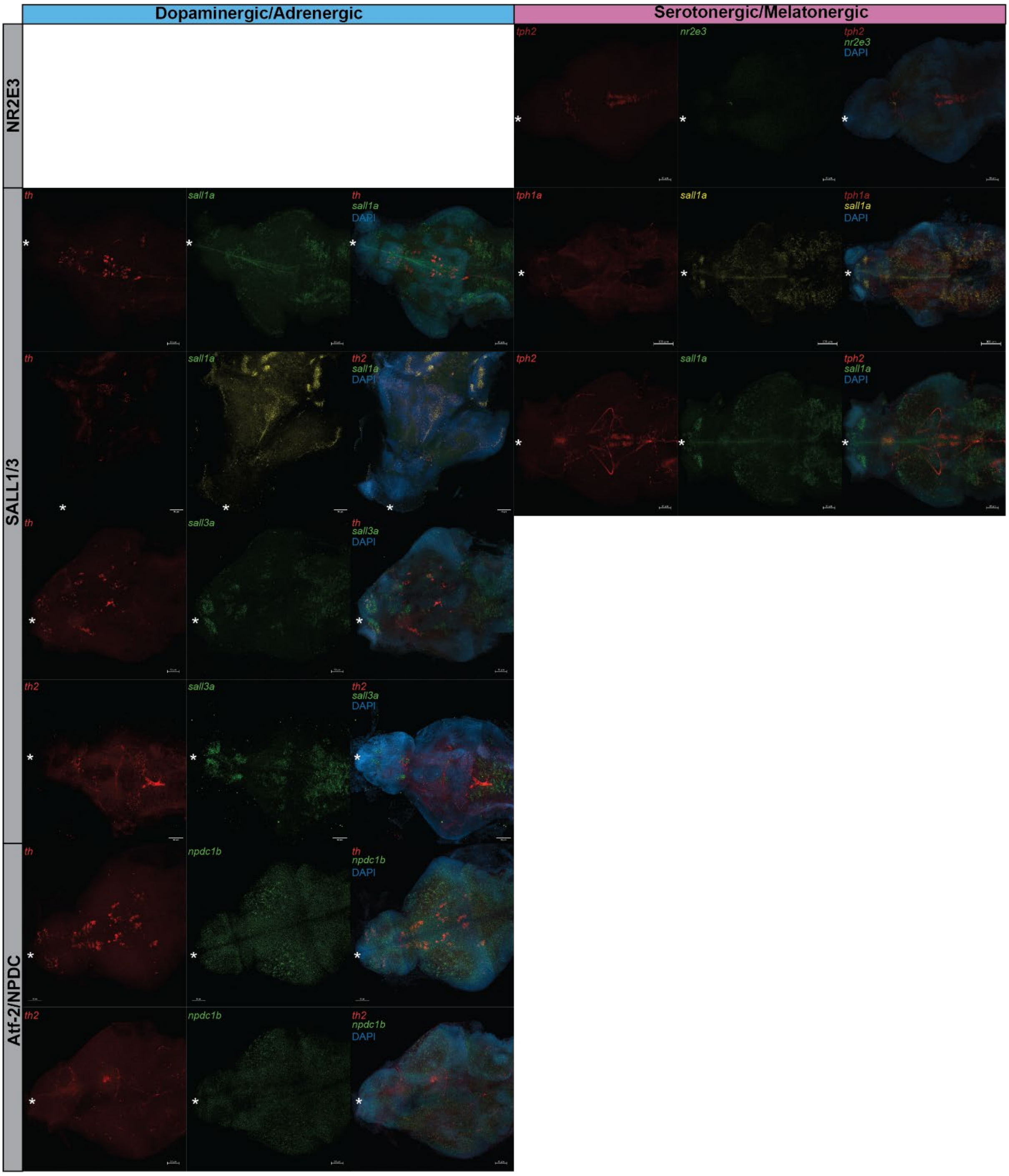

Dopaminergic/Adrenergic
Serotonergic/Melatonergic
NR2E3
SALL1/3
Atf-2/NPDC
tph2
nr2e3
tph2
nr2e3
DAPI
th
sall1a
th
sall1a
DAPI
tph1a
sall1a
tph1a
sall1a
DAPI
th
sall1a
th2
sall1a
DAPI
tph2
sall1a
tph2
sall1a
DAPI
th
sall3a
th
sall3a
DAPI
th2
sall3a
th2
sall3a
DAPI
th
npdc1b
th
npdc1b
DAPI
th2
npdc1b
th2
npdc1b
DAPI

**Figure S15: Panel of HCR images on *D. rerio* 5 days post fertilisation brains**

Images of *In Situ* hybridisation chain reaction staining of monoaminergic biosynthetic enzymes and transcription factor expression in 5-day old *Danio rerio* brains. Panels split showing expression of the enzyme and transcription factor independently as well as a combined image with DAPI staining. Enzymes shown in red. Transcription factors show in green or yellow. DAPI shown in blue. Gene names shown in top left corner of images. Orthogroup the transcription factors belong to is shown along the left. The monoamine the enzyme is linked to shown along the top. Gene and probe information in Table S5. White asterisk indicated the medial anterior point of the brain.

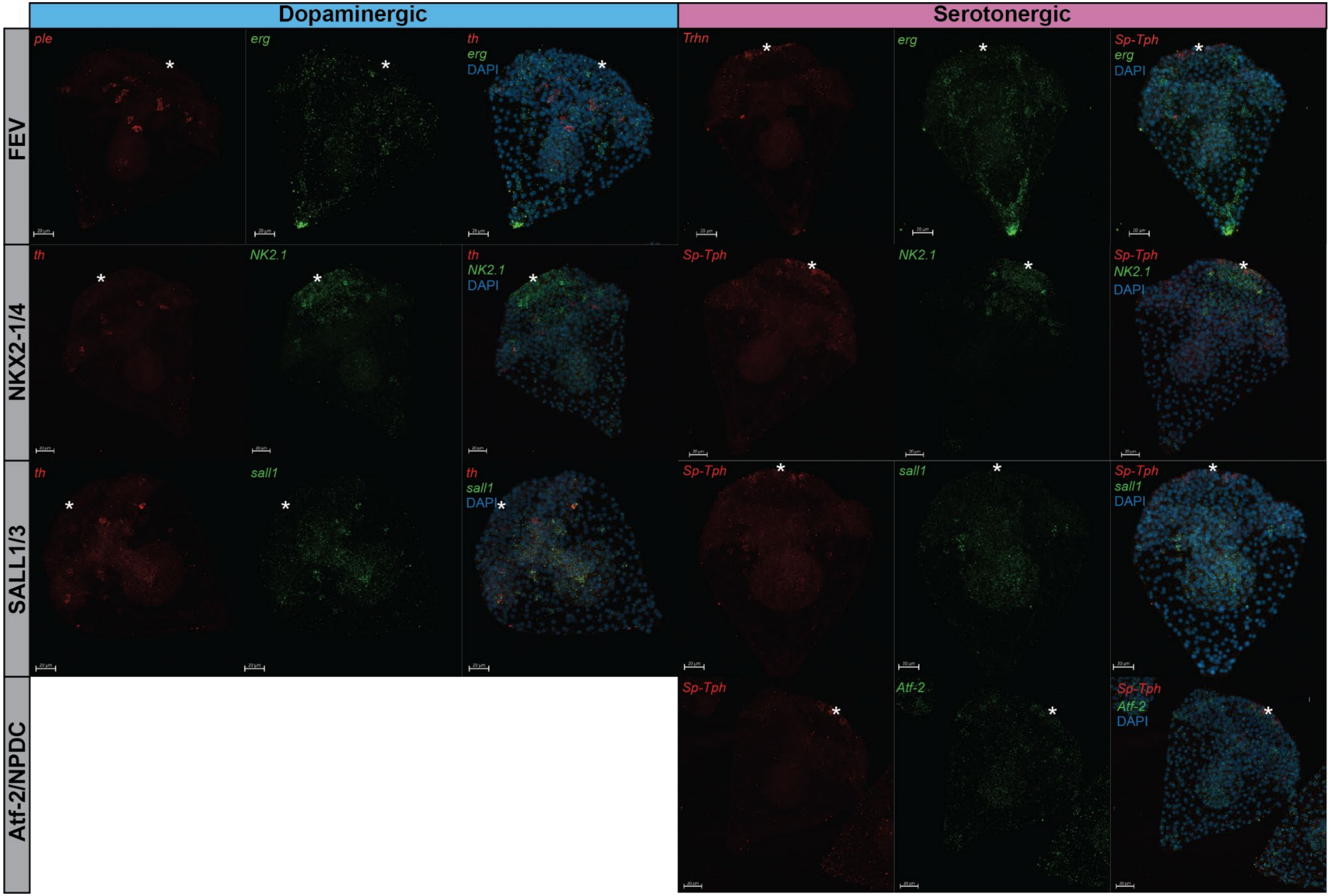

Dopaminergic
Serotonergic
FEV
NKX2-1/4
SALL1/3
Atf-2/NPDC
ple
erg
th
erg
DAPI
Trhn
erg
Sp-Tph
erg
DAPI
th
NK2.1
th
NK2.1
DAPI
Sp-Tph
NK2.1
Sp-Tph
NK2.1
DAPI
th
sall1
th
sall1
DAPI
Sp-Tph
sall1
Sp-Tph
sall1
DAPI
Sp-Tph
Atf-2
Sp-Tph
Atf-2
DAPI

**Figure S16: Panel of HCR images on *S. purpuratus* 5 days post fertilisation larvae**

Images of *In Situ* hybridisation chain reaction staining of monoaminergic biosynthetic enzymes and transcription factor expression in 3-day old *Strongylocentrotus purpuratus* larvae. Panels split showing expression of the enzyme and transcription factor independently as well as a combined image with DAPI staining. Enzymes shown in red. Transcription factors show in green or yellow. DAPI shown in blue. Gene names shown in top left corner of images. Orthogroup the transcription factors belong to is shown along the left. The monoamine the enzyme is linked to shown along the top. Gene and probe information in Table S5. White asterisk indicated the medial anterior point of the brain.

A)

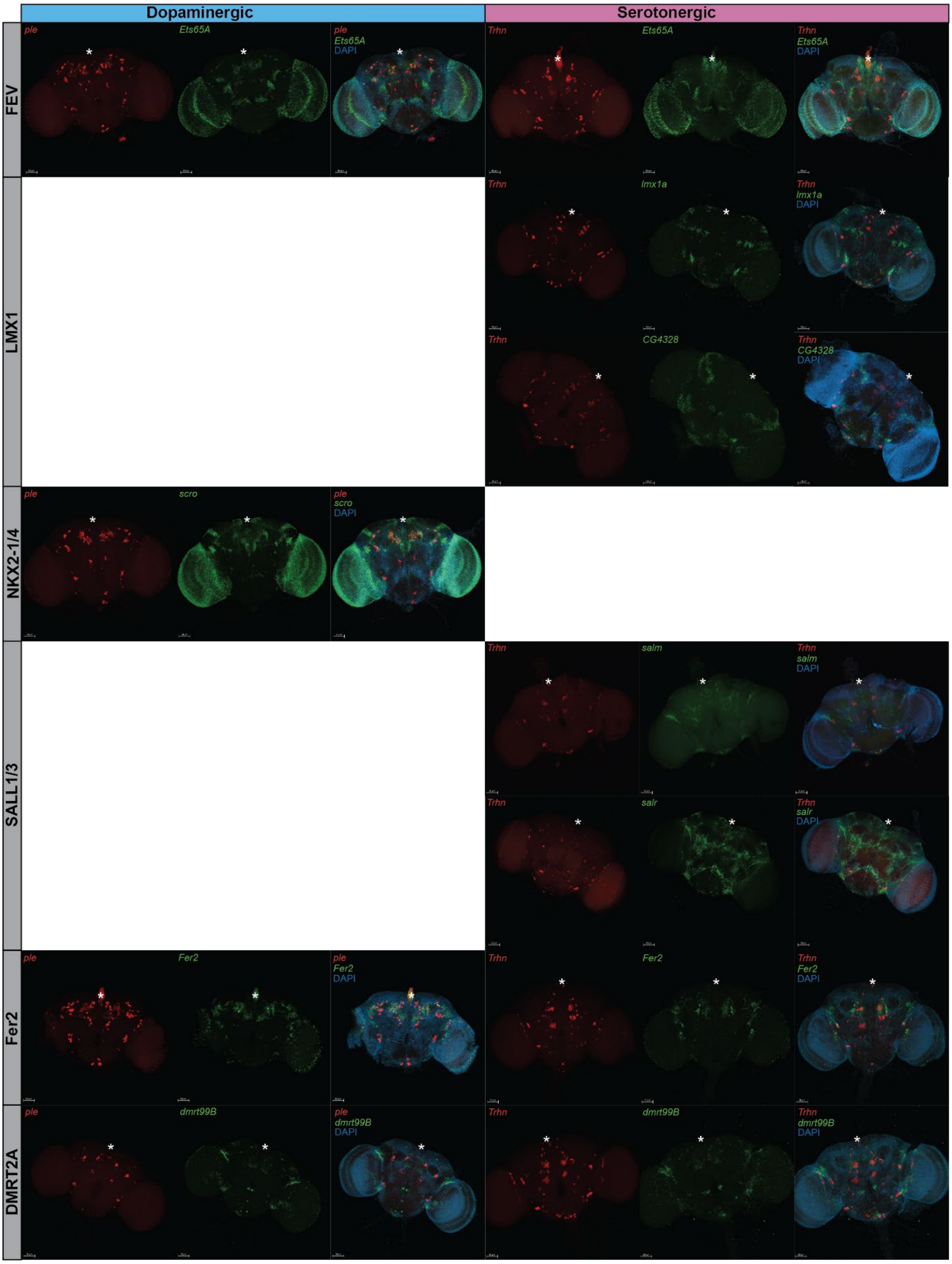
Dopaminergic
Serotonergic
FEV
LMX1
NKX2-1/4
SALL1/3
Fer2
DMRT2A
ple
Ets65A
ple Ets65A DAPI
Trhn
Ets65A
Trhn Ets65A DAPI
Trhn
lmx1a
Trhn lmx1a DAPI
Trhn
CG4328
Trhn CG4328 DAPI
ple
scro
ple scro DAPI
Trhn
salm
Trhn salm DAPI
Trhn
salr
Trhn salr DAPI
ple
Fer2
ple Fer2 DAPI
Trhn
Fer2
Trhn Fer2 DAPI
ple
dmrt99B
ple dmrt99B DAPI
Trhn
dmrt99B
Trhn dmrt99B DAPI

B)

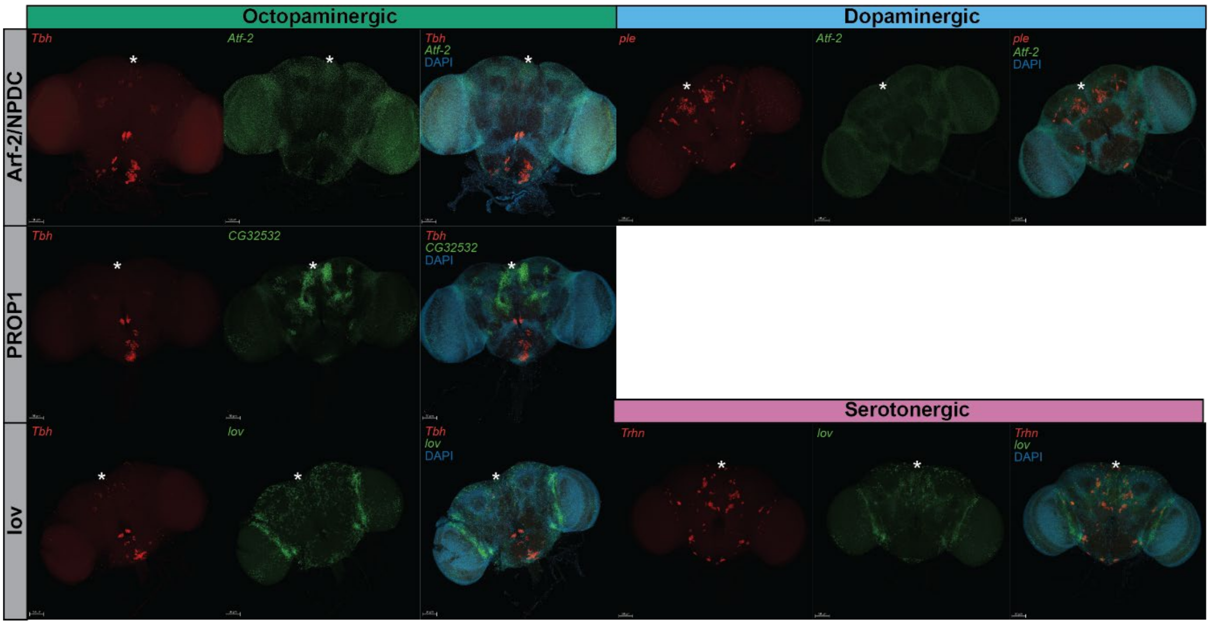


**Figure S17: Panel of HCR images on adult *D. melanogaster* brains**

Images of *In Situ* hybridisation chain reaction staining of monoaminergic biosynthetic enzymes and transcription factor expression in adult *Drosophila melanogaster* brains. Panels split showing expression of the enzyme and transcription factor independently as well as a combined image with DAPI staining. Enzymes shown in red. Transcription factors show in green or yellow. DAPI shown in blue. Gene names shown in top left corner of images. Orthogroup the transcription factors belong to is shown along the left. The monoamine the enzyme is linked to shown along the top. Gene and probe information in Table S5. White asterisk indicated the medial anterior point of the brain.

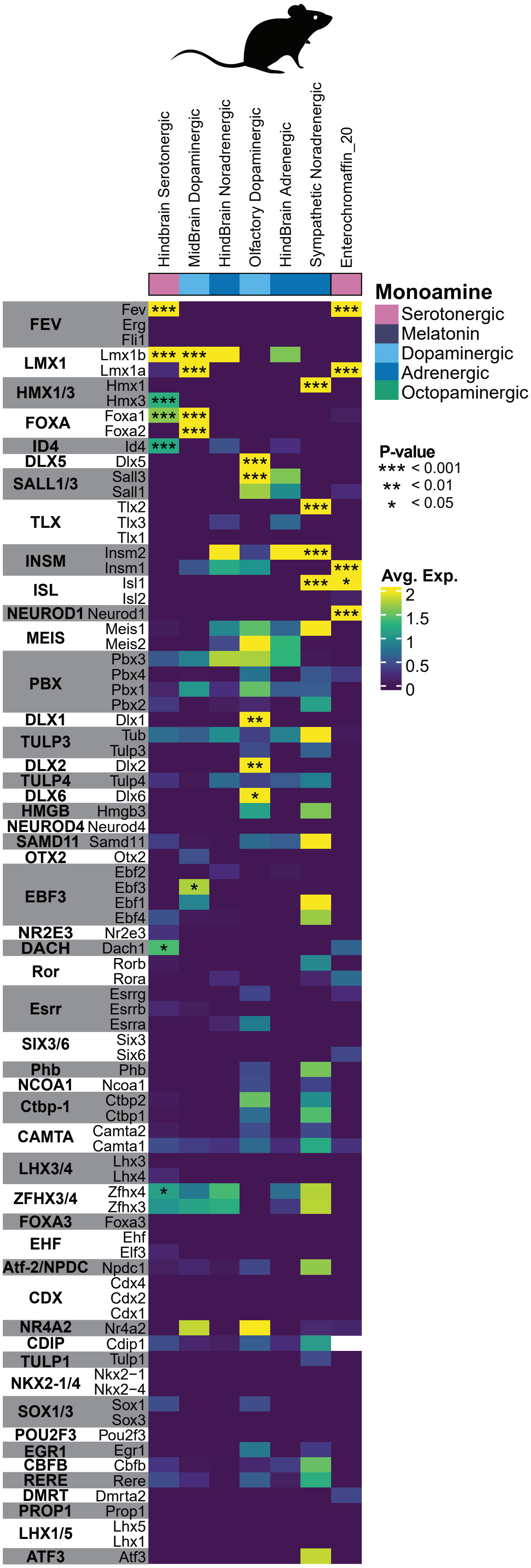

Hindbrain Serotonergic
MidBrain Dopaminergic
HindBrain Noradrenergic
Olfactory Dopaminergic
HindBrain Adrenergic
Sympathetic Noradrenergic
Enterochromaffin_20
Monoamine
Serotonergic
Melatonin
Dopaminergic
Adrenergic
Octopaminergic
P-value
*** < 0.001
** < 0.01
* < 0.05
Avg. Exp.
2
1.5
1
0.5
0
FEV
Fev
Erg
Fli1
LMX1
Lmx1b
Lmx1a
HMX1/3
Hmx1
Hmx3
FOXA
Foxa1
Foxa2
ID4
Id4
DLX5
Dlx5
SALL1/3
Sall3
Sall1
TLX
Tlx2
Tlx3
Tlx1
INSM
Insm2
Insm1
ISL
Isl1
Isl2
NEUROD1
Neurod1
MEIS
Meis1
Meis2
PBX
Pbx3
Pbx4
Pbx1
Pbx2
DLX1
Dlx1
TULP3
Tub
Tulp3
DLX2
Dlx2
TULP4
Tulp4
DLX6
Dlx6
HMGB
Hmgb3
NEUROD4
Neurod4
SAMD11
Samd11
OTX2
Otx2
EBF3
Ebf2
Ebf3
Ebf1
Ebf4
NR2E3
Nr2e3
DACH
Dach1
Ror
Rorb
Rora
Esrr
Esrrg
Esrrb
Esrra
SIX3/6
Six3
Six6
Phb
Phb
NCOA1
Ncoa1
Ctbp-1
Ctbp2
Ctbp1
CAMTA
Camta2
Camta1
LHX3/4
Lhx3
Lhx4
ZFHX3/4
Zfhx4
Zfhx3
FOXA3
Foxa3
EHF
Ehf
Elf3
Atf-2/NPDC
Npdc1
CDX
Cdx4
Cdx2
Cdx1
NR4A2
Nr4a2
CDIP
Cdip1
TULP1
Tulp1
NKX2-1/4
Nkx2−1
Nkx2−4
SOX1/3
Sox1
Sox3
POU2F3
Pou2f3
EGR1
Egr1
CBFB
Cbfb
RERE
Rere
DMRT
Dmrta2
PROP1
Prop1
LHX1/5
Lhx5
Lhx1
ATF3
Atf3

**Figure S18: *Mus musculus* Pseudobulk Expression Heatmap of Transcription Factors from Orthogroups shared across Species**

Heatmaps of transcription factor expression from *Mus musculus* pseudobulk analyses. Rows show individual homolog expression grouped in grey and white regions to indicate orthogroups. Only monoaminergic clusters shown with monoamine subtype colour coded along the top. Black boxes and labels group clusters from the same dataset. Significance of differential expression. indicated by *. Orthogroup names shown along the right. Expression is shown for all members of orthogroups recovered as significantly shared across species in one of the integrated monoaminergic clusters and in the unintegrated monoaminergic clusters of least two model organisms. Silhouette image from Phylopic.org: silhouette image by Daniel Jaron (*Mus musculus*).

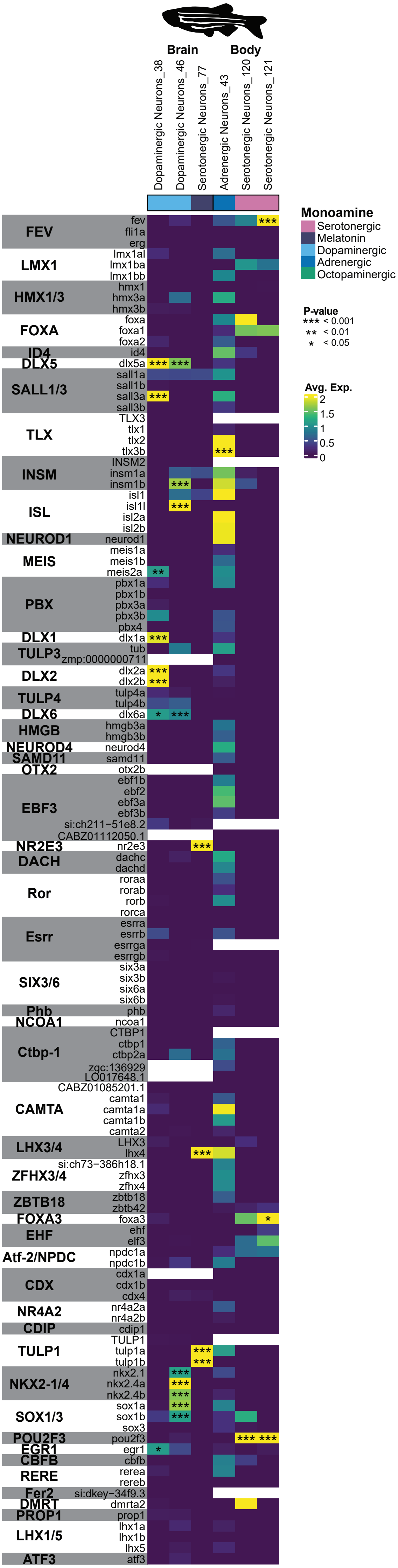

Brain
Body
Dopaminergic Neurons_38
Dopaminergic Neurons_46
Serotonergic Neurons_77
Adrenergic Neurons_43
Serotonergic Neurons_120
Serotonergic Neurons_121
Monoamine
Serotonergic
Melatonin
Dopaminergic
Adrenergic
Octopaminergic
P-value
*** < 0.001
** < 0.01
* < 0.05
Avg. Exp.
2
1.5
1
0.5
0
FEV
fev
fli1a
erg
LMX1
lmx1al
lmx1ba
lmx1bb
HMX1/3
hmx1
hmx3a
hmx3b
FOXA
foxa
foxa1
foxa2
ID4
id4
DLX5
dlx5a
SALL1/3
sall1a
sall1b
sall3a
sall3b
TLX
TLX3
tlx1
tlx2
tlx3b
INSM
INSM2
insm1a
insm1b
ISL
isl1
isl1l
isl2a
isl2b
NEUROD1
neurod1
MEIS
meis1a
meis1b
meis2a
PBX
pbx1a
pbx1b
pbx3a
pbx3b
pbx4
DLX1
dlx1a
TULP3
tub
zmp:0000000711
DLX2
dlx2a
dlx2b
TULP4
tulp4a
tulp4b
DLX6
dlx6a
HMGB
hmgb3a
hmgb3b
NEUROD4
neurod4
SAMD11
samd11
OTX2
otx2b
EBF3
ebf1b
ebf2
ebf3a
ebf3b
si:ch211−51e8.2
CABZ01112050.1
NR2E3
nr2e3
DACH
dachc
dachd
Ror
roraa
rorab
rorb
rorca
Esrr
esrra
esrrb
esrrga
esrrgb
SIX3/6
six3a
six3b
six6a
six6b
Phb
phb
NCOA1
ncoa1
Ctbp-1
CTBP1
ctbp1
ctbp2a
zgc:136929
LO017648.1
CABZ01085201.1
CAMTA
camta1
camta1a
camta1b
camta2
LHX3/4
LHX3
lhx4
ZFHX3/4
si:ch73−386h18.1
zfhx3
zfhx4
ZBTB18
zbtb18
zbtb42
FOXA3
foxa3
EHF
ehf
elf3
Atf-2/NPDC
npdc1a
npdc1b
CDX
cdx1a
cdx1b
cdx4
NR4A2
nr4a2a
nr4a2b
CDIP
cdip1
TULP1
TULP1
tulp1a
tulp1b
NKX2-1/4
nkx2.1
nkx2.4a
nkx2.4b
SOX1/3
sox1a
sox1b
sox3
POU2F3
pou2f3
EGR1
egr1
CBFB
cbfb
RERE
rerea
rereb
Fer2
si:dkey−34f9.3
DMRT
dmrta2
PROP1
prop1
LHX1/5
lhx1a
lhx1b
lhx5
ATF3
atf3

**Figure S18: *Danio rerio* Pseudobulk Expression Heatmap of Transcription Factors from Orthogroups shared across Species**

Heatmaps of transcription factor expression from *Danio rerio* pseudobulk analyses. Rows show individual homolog expression grouped in grey and white regions to indicate orthogroups. Only monoaminergic clusters shown with monoamine subtype colour coded along the top. Black boxes and labels group clusters from the same dataset. Significance of differential expression. indicated by *. Orthogroup names shown along the right. Expression is shown for all members of orthogroups recovered as significantly shared across species in one of the integrated monoaminergic clusters and in the unintegrated monoaminergic clusters of least two model organisms. Silhouette image from Phylopic.org: silhouette image by Jake Warner (*Danio rerio*).

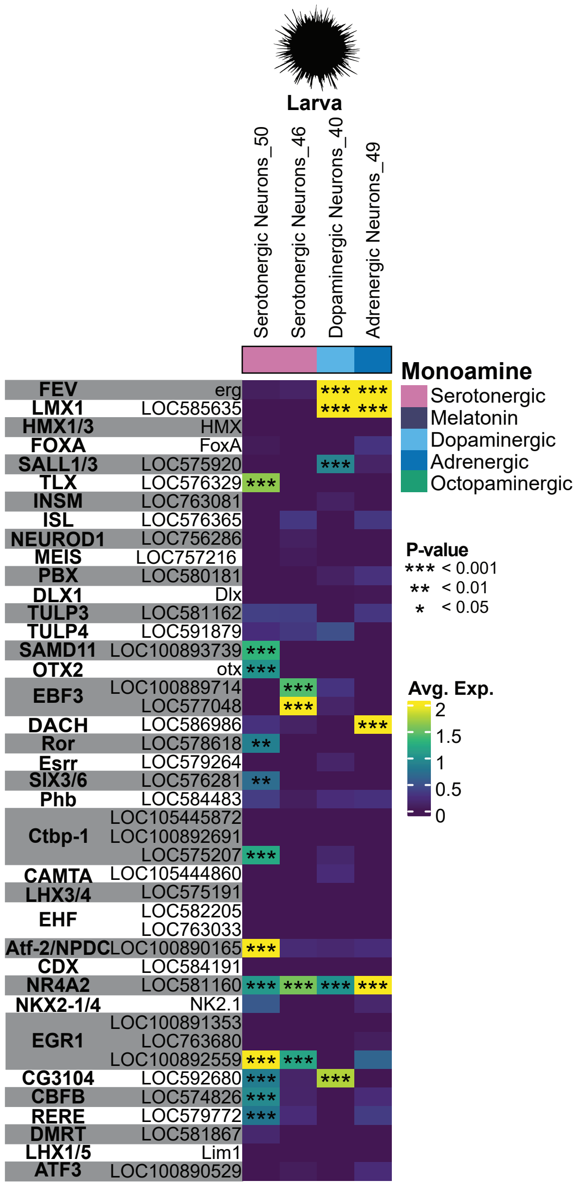

Larva
Serotonergic Neurons_50
Serotonergic Neurons_46
Dopaminergic Neurons_40
Adrenergic Neurons_49
Monoamine
Serotonergic
Melatonin
Dopaminergic
Adrenergic
Octopaminergic
P-value
*** < 0.001
** < 0.01
* < 0.05
Avg. Exp.
2
1.5
1
0.5
0
FEV erg
LMX1 LOC585635
HMX1/3 HMX
FOXA FoxA
SALL1/3 LOC575920
TLX LOC576329
INSM LOC763081
ISL LOC576365
NEUROD1 LOC756286
MEIS LOC757216
PBX LOC580181
DLX1 Dlx
TULP3 LOC581162
TULP4 LOC591879
SAMD11 LOC100893739
OTX2 otx
EBF3 LOC100889714
LOC577048
DACH LOC586986
Ror LOC578618
Esrr LOC579264
SIX3/6 LOC576281
Phb LOC584483
Ctbp-1 LOC105445872
LOC100892691
LOC575207
CAMTA LOC105444860
LHX3/4 LOC575191
EHF LOC582205
LOC763033
Atf-2/NPDC LOC100890165
CDX LOC584191
NR4A2 LOC581160
NKX2-1/4 NK2.1
EGR1 LOC100891353
LOC763680
LOC100892559
CG3104 LOC592680
CBFB LOC574826
RERE LOC579772
DMRT LOC581867
LHX1/5 Lim1
ATF3 LOC100890529

**Figure S20: *Strongylocentrotus purpuratus* Expression Heatmap of Transcription Factors from Orthogroups shared across Species**

Heatmaps of transcription factor expression from *Strongylocentrotus purpuratus* non-recoded analyses. Rows show individual homolog expression grouped in grey and white regions to indicate orthogroups. Only monoaminergic clusters shown with monoamine subtype colour coded along the top. Black boxes and labels group clusters from the same dataset. Significance of differential expression. indicated by *. Orthogroup names shown along the right. Expression is shown for all members of orthogroups recovered as significantly shared across species in one of the integrated monoaminergic clusters and in the unintegrated monoaminergic clusters of least two model organisms. Silhouette image from Phylopic.org: silhouette image by Christoph Schomburg (*Strongylocentrotus purpuratus*).

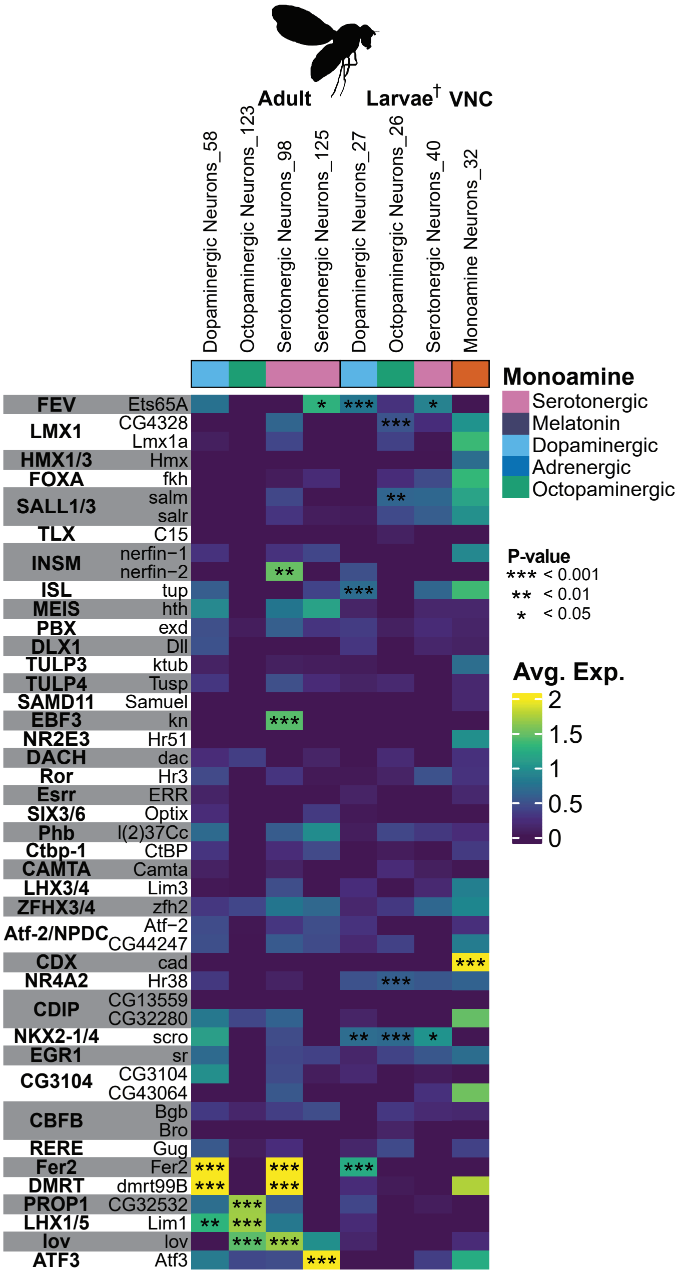

Adult
Larvae† VNC
Dopaminergic Neurons_58
Octopaminergic Neurons_123
Serotonergic Neurons_98
Serotonergic Neurons_125
Dopaminergic Neurons_27
Octopaminergic Neurons_26
Serotonergic Neurons_40
Monoamine Neurons_32
Monoamine
Serotonergic
Melatonin
Dopaminergic
Adrenergic
Octopaminergic
P-value
*** < 0.001
** < 0.01
* < 0.05
Avg. Exp.
2
1.5
1
0.5
0
FEV Ets65A
LMX1 CG4328 Lmx1a
HMX1/3 Hmx
FOXA fkh
SALL1/3 salm salr
TLX C15
INSM nerfin−1 nerfin−2
ISL tup
MEIS hth
PBX exd
DLX1 Dll
TULP3 ktub
TULP4 Tusp
SAMD11 Samuel
EBF3 kn
NR2E3 Hr51
DACH dac
Ror Hr3
Esrr ERR
SIX3/6 Optix
Phb l(2)37Cc
Ctbp-1 CtBP
CAMTA Camta
LHX3/4 Lim3
ZFHX3/4 zfh2
Atf-2/NPDC Atf−2 CG44247
CDX cad
NR4A2 Hr38
CDIP CG13559 CG32280
NKX2-1/4 scro
EGR1 sr
CG3104 CG3104 CG43064
CBFB Bgb Bro
RERE Gug
Fer2 Fer2
DMRT dmrt99B
PROP1 CG32532
LHX1/5 Lim1
lov lov
ATF3 Atf3

**Figure S21: *Drosophila melanogaster* Pseudobulk Expression Heatmap of Transcription Factors from Orthogroups shared across Species**

Heatmaps of transcription factor expression from *Drosophila melanogaster* pseudobulk analyses. Non-peudobulked datasets indicated with †. Rows show individual homolog expression grouped in grey and white regions to indicate orthogroups. Only monoaminergic clusters shown with monoamine subtype colour coded along the top. Black boxes and labels group clusters from the same dataset. Significance of differential expression. indicated by *. Orthogroup names shown along the right. Expression is shown for all members of orthogroups recovered as significantly shared across species in one of the integrated monoaminergic clusters and in the unintegrated monoaminergic clusters of least two model organisms. Silhouette image from Phylopic.org: silhouette image by Ramiro Morales-Hojas (*Drosophila americana* as *Drosophila melanogaster*).

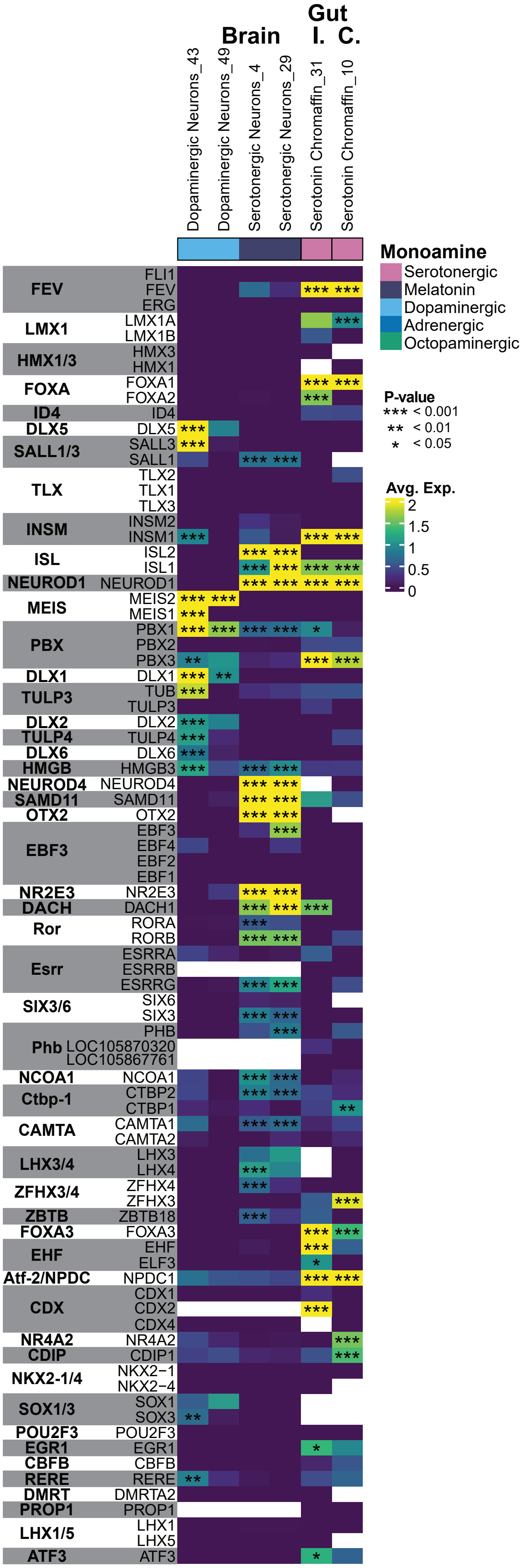

Brain
Gut
I. C.
Dopaminergic Neurons_43
Dopaminergic Neurons_49
Serotonergic Neurons_4
Serotonergic Neurons_29
Serotonin Chromaffin_31
Serotonin Chromaffin_10
Monoamine
Serotonergic
Melatonin
Dopaminergic
Adrenergic
Octopaminergic
P-value
*** < 0.001
** < 0.01
* < 0.05
Avg. Exp.
2
1.5
1
0.5
0
FEV FLI1 FEV ERG
LMX1 LMX1A LMX1B
HMX1/3 HMX3 HMX1
FOXA FOXA1 FOXA2
ID4 ID4
DLX5 DLX5
SALL1/3 SALL3 SALL1
TLX TLX2 TLX1 TLX3
INSM INSM2 INSM1
ISL ISL2 ISL1
NEUROD1 NEUROD1
MEIS MEIS2 MEIS1
PBX PBX1 PBX2 PBX3
DLX1 DLX1
TULP3 TUB TULP3
DLX2 DLX2
TULP4 TULP4
DLX6 DLX6
HMGB HMGB3
NEUROD4 NEUROD4
SAMD11 SAMD11
OTX2 OTX2
EBF3 EBF3 EBF4 EBF2 EBF1
NR2E3 NR2E3
DACH DACH1
Ror RORA RORB
Esrr ESRRA ESRRB ESRRG
SIX3/6 SIX6 SIX3
Phb PHB LOC105870320 LOC105867761
NCOA1 NCOA1
Ctbp-1 CTBP2 CTBP1
CAMTA CAMTA1 CAMTA2
LHX3/4 LHX3 LHX4
ZFHX3/4 ZFHX4 ZFHX3
ZBTB ZBTB18
FOXA3 FOXA3
EHF EHF ELF3
Atf-2/NPDC NPDC1
CDX CDX1 CDX2 CDX4
NR4A2 NR4A2
CDIP CDIP1
NKX2-1/4 NKX2−1 NKX2−4
SOX1/3 SOX1 SOX3
POU2F3 POU2F3
EGR1 EGR1
CBFB CBFB
RERE RERE
DMRT DMRTA2
PROP1 PROP1
LHX1/5 LHX1 LHX5
ATF3 ATF3

**Figure S22: *Microcebus murinus* Expression Heatmap of Transcription Factors from Orthogroups shared across Species**

Heatmaps of transcription factor expression from *Microcebus murinus* non-recoded analyses. Rows show individual homolog expression grouped in grey and white regions to indicate orthogroups. Only monoaminergic clusters shown with monoamine subtype colour coded along the top. Black boxes and labels group clusters from the same dataset. Significance of differential expression. indicated by *. Orthogroup names shown along the right. Expression is shown for all members of orthogroups recovered as significantly shared across species in one of the integrated monoaminergic clusters and in the unintegrated monoaminergic clusters of least two model organisms.

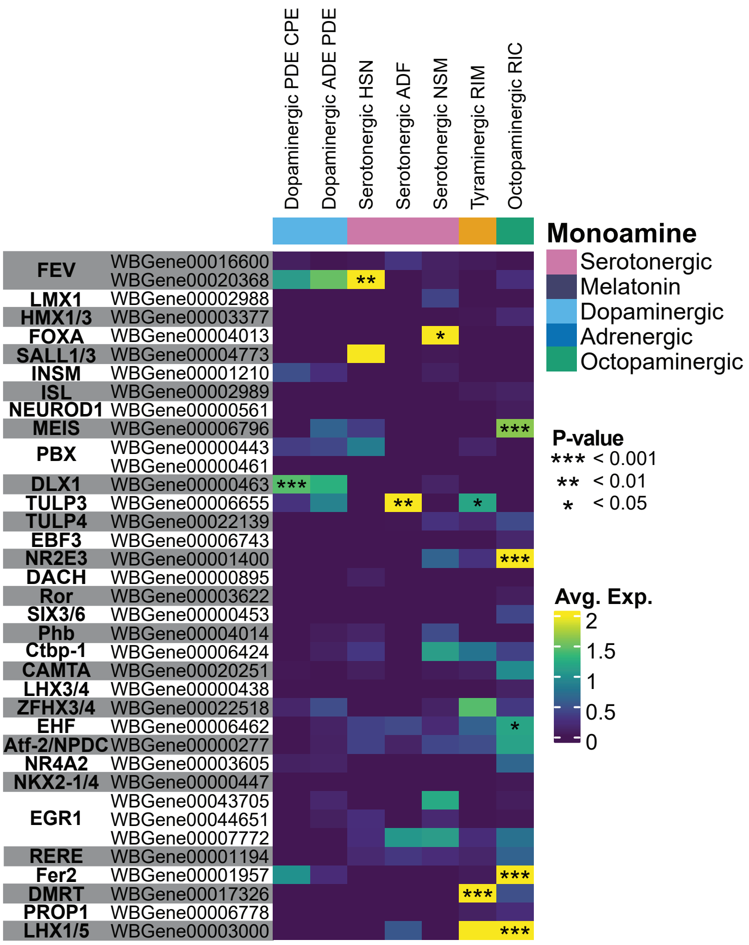

Dopaminergic PDE CPE
Dopaminergic ADE PDE
Serotonergic HSN
Serotonergic ADF
Serotonergic NSM
Tyraminergic RIM
Octopaminergic RIC
Monoamine
Serotonergic
Melatonin
Dopaminergic
Adrenergic
Octopaminergic
FEV WBGene00016600
WBGene00020368
LMX1 WBGene00002988
HMX1/3 WBGene00003377
FOXA WBGene00004013
SALL1/3 WBGene00004773
INSM WBGene00001210
ISL WBGene00002989
NEUROD1 WBGene00000561
MEIS WBGene00006796
PBX WBGene00000443
WBGene00000461
DLX1 WBGene00000463
TULP3 WBGene00006655
TULP4 WBGene00022139
EBF3 WBGene00006743
NR2E3 WBGene00001400
DACH WBGene00000895
Ror WBGene00003622
SIX3/6 WBGene00000453
Phb WBGene00004014
Ctbp-1 WBGene00006424
CAMTA WBGene00020251
LHX3/4 WBGene00000438
ZFHX3/4 WBGene00022518
EHF WBGene00006462
Atf-2/NPDC WBGene00000277
NR4A2 WBGene00003605
NKX2-1/4 WBGene00000447
EGR1 WBGene00043705
WBGene00044651
WBGene00007772
RERE WBGene00001194
Fer2 WBGene00001957
DMRT WBGene00017326
PROP1 WBGene00006778
LHX1/5 WBGene00003000
P-value
*** < 0.001
** < 0.01
* < 0.05
Avg. Exp.
2
1.5
1
0.5
0

**Figure S23: *Caenorhabditis elegans* Pseudobulk Expression Heatmap of Transcription Factors from Orthogroups shared across Species**

Heatmaps of transcription factor expression from *Caenorhabditis elegans* pseudobulk analyses. Rows show individual homolog expression grouped in grey and white regions to indicate orthogroups. Only monoaminergic clusters shown with monoamine subtype colour coded along the top. Black boxes and labels group clusters from the same dataset. Significance of differential expression. indicated by *. Orthogroup names shown along the right. Expression is shown for all members of orthogroups recovered as significantly shared across species in one of the integrated monoaminergic clusters and in the unintegrated monoaminergic clusters of least two model organisms.

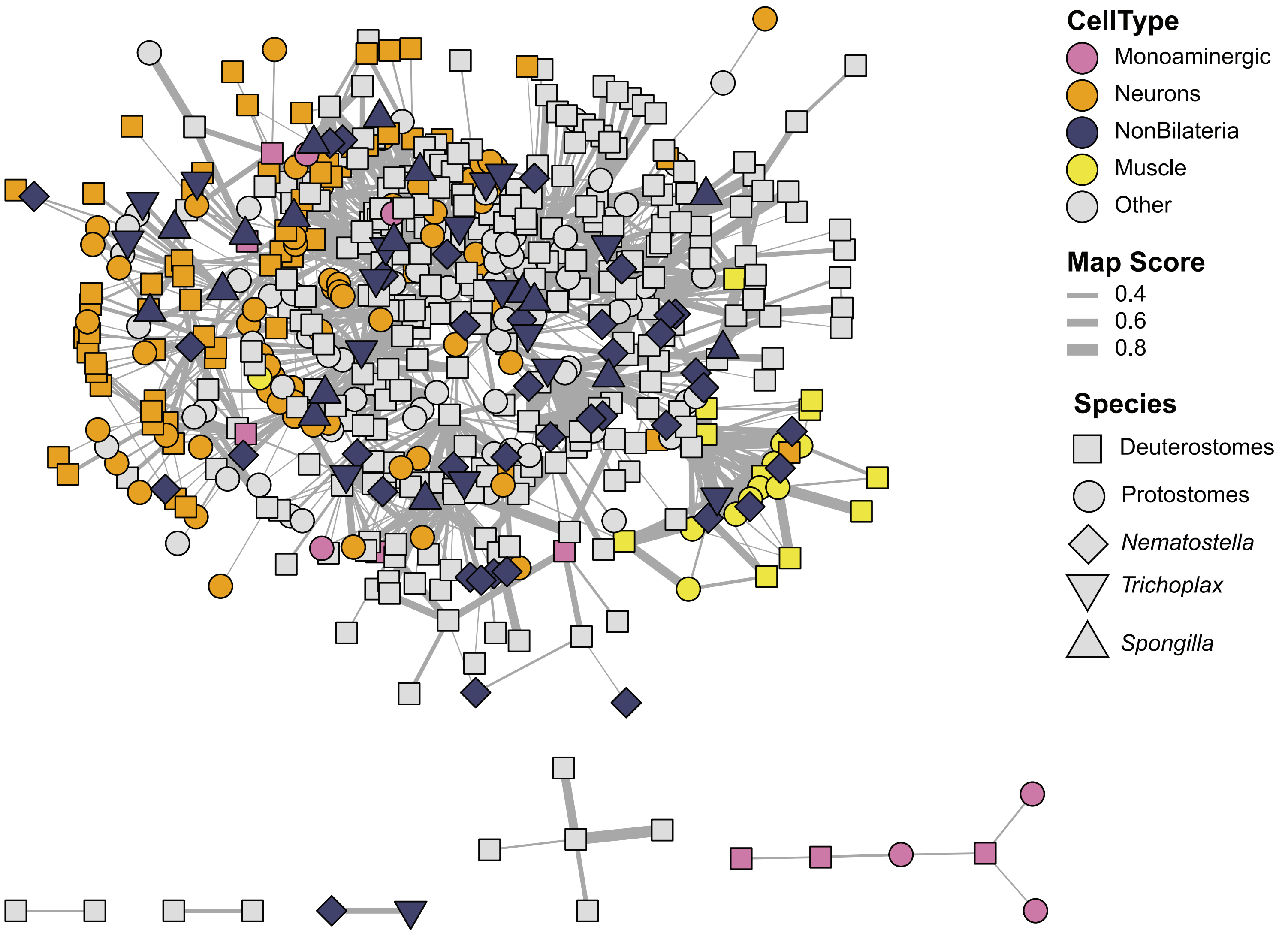
CellType
Monoaminergic
Neurons
NonBilateria
Muscle
Other
Map Score
0.4
0.6
0.8
Species
Deuterostomes
Protostomes
*Nematostella*
*Trichoplax*
*Spongilla*

**Figure S24: Monoaminergic Neurons do not share Transcriptional Profiles with Non-bilaterians**

Network of SAMap mapping scores between individual cell clusters from the three non-bilaterian datasets (*Nematostella vectensis*, *Trichoplax adhaerens* and *Spongilla lacustrus*) and the six species, model species dataset. Nodes represent cell clusters and edges represent SAMap mapping scores. Edge thickness denoted the mapping strength, scores below 0.2 are omitted. Nodes are colour coded by broad cell type group.
