## Supplementary Table Info for "Monoaminergic neurons share transcriptional identity across Bilaterian animals"

**Table S1: Known Monoaminergic Associated Transcription Factors**

Sources[1–43]

**Table S2: Single-Cell RNAseq and Proteome Data Sources**

**Table S3: Bilaterian Brain SAMap Homolog Alignment Scores**

**Table S4: List of TF Orthogroups**

**Table S5: HCR Probe Sequences**

**Table S6: Software Used**

**Table S7: Reagents Used**
